# Deciphering and Improving Human Homogentisate 1,2-Dioxygenase Function Through Knowledge Gaining Directed Evolution: Implications for Alkaptonuria

**DOI:** 10.64898/2026.06.02.729205

**Authors:** Sien Lequeue, Jessie Neuckermans, Liesbeth Desmet, Nina S. Salvi, Tamara Vanhaecke, Ulrich Schwaneberg, Joery De Kock

**Affiliations:** Vrije Universiteit Brussel, In Vitro Toxicology and Dermato-Cosmetology (IVTD), Liver Therapy & Evolution Team, Faculty of Medicine and Pharmacy, 1090 Brussels, Belgium; Vrije Universiteit Brussel, In Vitro Toxicology and Dermato-Cosmetology (IVTD), In Vitro Liver Disease Modelling Team, Faculty of Medicine and Pharmacy, 1090 Brussels, Belgium; RWTH Aachen University, Lehrstuhl für Biotechnologie, Worringerweg 3, 52074 Aachen, Germany

**Keywords:** KnowVolution, homogentisate 1, 2-dioxygenase, alkaptonuria, genotype-phenotype correlations, protein structure-function, directed evolution, *Escherichia coli*

## Abstract

Human homogentisate 1,2-dioxygenase (HGD) catalyses the oxidative cleavage of homogentisic acid (HGA) to maleylacetoacetate (MAA), a key step in tyrosine degradation. Loss of HGD activity causes alkaptonuria (AKU), a rare inherited metabolic disorder characterized by toxic HGA accumulation. Current therapy with nitisinone lowers HGA levels but does not restore HGD function, motivating further investigation of HGD structure-function relationships. In this study, we applied the Knowledge Gaining Directed Evolution (KnowVolution) strategy to investigate how amino acid substitutions influence catalytic activity and structural integrity of human HGD. Catalytic activity was evaluated in *Escherichia coli* using an assay quantifying MAA formation over time. Across four KnowVolution phases, multiple substitutions were identified that modulated catalytic activity while preserving enzyme function. Notably, none of the influential substitutions were located within the catalytic pocket; instead, they occurred predominantly at surface-exposed or structural positions. Structural mapping, interface analysis, and computational stability predictions indicated that some substitutions contribute to hexamer stabilization, whereas others likely alter activity through indirect, non-catalytic mechanisms involving pocket remodelling. Combined substitutions showed non-additive effects that were either cooperative or antagonistic, demonstrating that their impact could not be predicted from individual contributions. Tunnel and pocket analyses showed that N31S, S54D and D86H produced a more compact hexamer, whereas a Q354P+P359E double mutant reduced catalytic pocket solvent accessibility and volume, supporting the observed activity differences. Overall, these findings demonstrate that HGD activity can be modulated by substitutions outside the catalytic pocket, providing new insight into HGD function and genotype-phenotype relationships underlying AKU.

## 1. Introduction

Human homogentisate 1,2-dioxygenase (HGD; EC 1.13.11.5) catalyses the oxidative cleavage of homogentisic acid (HGA) to maleylacetoacetate (MAA) (**Fig. 1a**), a critical step in the tyrosine degradation pathway. Loss of HGD function disrupts this reaction and causes alkaptonuria (AKU; OMIM #203500), a rare autosomal recessive metabolic disorder [1]. AKU is caused by homozygous or compound heterozygous mutations within the HGD gene, a single-copy gene located on chromosome 3q21-q23 comprising 14 exons. So far, 259 unique pathogenic DNA variants have been identified in 748 patients (January 2026), with the majority being missense mutations [2–6]. Although some variants retain partial enzyme activity, this is generally insufficient to prevent HGA accumulation. Biallelic pathogenic variants in HGD lead to progressive HGA buildup, which undergoes oxidation and polymerization into a melanin-like ochronotic pigment. The consequent pigment deposition in connective body tissues produces the hallmark clinical features of AKU, including dark urine and bluish-black discoloration of cartilage, sclera, and skin. Serious complications can also arise, such as aortic-valve stenosis, vascular calcification, nephrolithiasis, and an early-onset, debilitating arthropathy that markedly reduces quality of life [1,7,8]. Current therapies, such as nitisinone (NTBC), reduce HGA levels but do not restore HGD enzymatic function and can cause hypertyrosinemia, which may lead to severe side effects including leukopenia, thrombocytopenia, and ocular complications. These limitations underscore the need to better understand the structural and catalytic properties of HGD in relation to enzyme function, as a basis for developing alternative therapeutic strategies aimed at restoring HGD activity [9–11].

**Figure 1.**
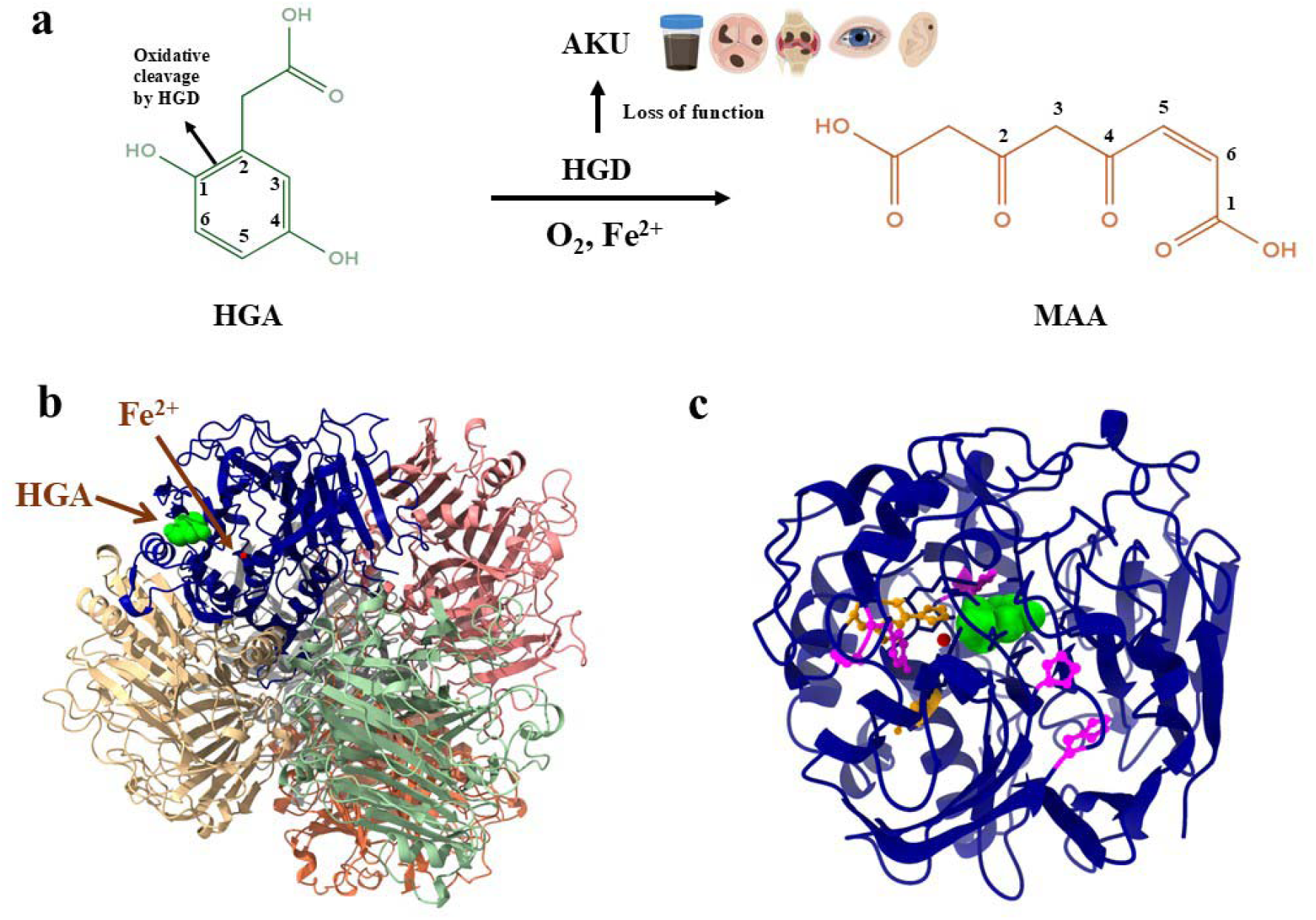
Structure and catalytic mechanism of human HGD. **(a)** Chemical conversion of HGA to MAA, catalysed by HGD (structures obtained from ChemSpider). Deficiency of HGD leads to AKU, illustrated with BioRender cartoons (https://BioRender.com/3nh5fuv), showing hallmark clinical features: dark urine, pigment deposition in heart valves and knee joints resulting in arthritis, and bluish-black discoloration of sclera and ear cartilage. **(b)** Hexameric structure of HGD (cartoon representation) with HGA (green) and Fe²□ (red) shown as spheres. The hexamer is arranged as a dimer of trimers, highlighting the overall quaternary assembly. **(c)** Key residues involved in the catalytic process of HGD. A single monomer is shown for clarity. Residues coordinating the Fe²□ ion (His335, Glu341, His371) are shown in orange as ball-and-stick representations, while residues interacting with the substrate HGA (His292, His365, Pro295, Pro332, Tyr333) are shown in magenta as ball-and-stick representations. His292 forms a hydrogen bond with the C4 hydroxyl of HGA, His365 participates in proton transfer during catalysis, and Pro295, Pro332, and Tyr333 form a solvent-accessible pocket that stabilizes substrate positioning. The substrate HGA is shown in green, and the Fe^2+^ ion is shown in red.

At the molecular level, the HGD enzyme forms a hexamer composed of six protomers arranged as a dimer of trimers (**Fig. 1b**). Each protomer consists of 445 amino acids (AAs) divided into an N-terminal domain of approximately 280 residues and a C-terminal domain of about 140 residues. The active site is located in the C-terminal domain near the interface between protomers, while the N-terminal domain forms a compact core that stabilizes the overall globular fold of the protein. The C-terminal domain has lower secondary structure content compared to the N-terminal domain, providing the flexibility needed to accommodate and position the substrate for catalysis [12–14]. Within the active site, a mononuclear non-heme Fe^2+^ ion is coordinated by His335, Glu341, and His371, with additional residues contributing to substrate binding. His292 forms a hydrogen bond with the C4 hydroxyl of HGA, stabilizing its orientation, and His365, hydrogen-bonded to Glu341, facilitates deprotonation and proton transfer during the reaction. Pro295, Pro332, and Tyr333 create a solvent-accessible pocket that supports substrate positioning. Catalysis begins with the carboxylate of HGA chelating the iron and deprotonation of the C2 hydroxyl group. Molecular dioxygen then coordinates to the Fe²⁺ center and attacks the aromatic ring specifically at the C1-C2 bond (**Fig. 1a**), which is activated by the C2 hydroxyl coordination. This results in insertion of oxygen between C1 and C2, leading to oxidative cleavage and opening of the aromatic ring *via* a peroxo-bridged intermediate. The intermediate rearranges into a transient lactone, which is then opened by nucleophilic attack of an iron-bound hydroxide. Subsequent tautomerization produces MAA, completing the irreversible oxidative cleavage and resetting the active site for the next catalytic cycle [12,13,15–18]. **Fig. 1c** highlights the key residues involved in this catalytic process (His335, Glu341, His371, His292, His365, Pro295, Pro332, and Tyr333) within the 3D structure of a single HGD protomer.

Because this enzymatic reaction depends on the precise quaternary assembly and active-site configuration of HGD, even subtle AA substitutions can destabilize the enzyme or impair catalysis, providing a molecular basis for the variable phenotypic outcomes observed in AKU. The folding of individual protomers and their association into the hexameric structure rely heavily on noncovalent interactions among AAs, meaning that missense variants can have severe consequences. Hydrophilic surface residues are particularly important for maintaining protein solubility, core residues stabilize the tertiary fold, and active-site residues are critical for catalytic function [14,19]. As a result, substitutions at these critical residues can not only compromise HGD’s structural integrity but also alter its enzymatic efficiency, providing a molecular explanation for the variable severity observed in AKU patients. However, despite extensive characterization of HGD variants, residual enzyme activity does not always predict disease severity, and most previous studies have been observational [3,6,14,20–22].

To address this, the Knowledge Gaining Directed Evolution (KnowVolution) strategy was employed, as previously reported to improve for instance arginine deiminase close to physiological conditions [23–26]. A full campaign was performed to systematically generate, test, and optimize human HGD variants under physiologically relevant conditions, providing a controlled approach to identify substitutions that improve enzyme stability and catalytic efficiency, while also gaining insights into structure-function relationships that could have added value for precision medicine approaches in AKU [14,27]. A KnowVolution campaign combines random and focused mutagenesis with computational analysis in four phases: (I) identifying potentially influential AA positions, (II) determining which substitutions improve function, (III) assessing whether these changes interact cooperatively based on structural data, and (IV) recombining the best substitutions guided by stability predictions to maximize improvements. Compared to traditional directed evolution, KnowVolution requires less screening efforts while providing detailed insight into the molecular determinants of enzyme activity [27]. Functional screening of the resulting HGD variants was performed by using a previously established high-throughput screening (HTS) assay in *Escherichia coli (E. coli)*, adapted from a shake-flask-based format and validated in a 96-well plate format ensuring a robust assessment of HGD’s enzyme activity [28]. Altogether, by linking sequence variants to functional outcomes, this strategy can shed light on the molecular basis of HGD deficiency, contribute to a more nuanced understanding of AKU phenotype variability, and support future precision medicine approaches.

## 2. Materials and methods

### 2.1 Materials

All chemicals used were of analytical reagent grade or higher and sourced from Sigma-Aldrich Chemie GmbH (Schnelldorf, Germany). Restriction enzymes, PCR master mix reagents, and T4 DNA ligase were obtained from Thermo Fisher Scientific (Waltham, MA, USA). All PCR reactions were performed using a Bio-Rad iCycler thermal cycler (Bio-Rad Laboratories, Hercules, CA, USA) with thin-walled PCR tubes (Sapphire PCR tube, 0.2 mL, polypropyleen; Greiner Bio-One, Kremsmünster, Austria). Plasmid extraction and PCR purification kits were purchased from Sigma-Aldrich (Hamburg, Germany). DNA concentrations were measured with a NanoDrop spectrophotometer (Thermo Fisher Scientific) and optical density at 600 nm (OD_600_) was determined using an Eppendorf BioPhotometer D30 (Hamburg, Germany). HGD protein expression was carried out in 96-well conical-bottom microtiter plates (Greiner Bio-One GmbH, Kremsmünster, Austria) and incubated in an Eppendorf New Brunswick™ Innova 42R shaker. Absorbance measurements for MAA assays were performed using a VICTOR 3 1420 Multilabel Counter plate reader (PerkinElmer, Waltham, MA, USA) with Nunc™ MicroWell™ 96-well microplates (Thermo Fisher Scientific).

### 2.2 Bacterial strains, plasmids, gene cloning and expression vector constructs

The cDNA of wildtype (WT) human HGD (NM_000187.3), codon-optimized for *E. coli*, was obtained from GeneArt (Thermo Scientific) in standard pMA-based cloning plasmids (see **Supplementary Data 1**). The insert was subcloned into the multiple cloning site of pET42b(+) expression vectors (Novagen, Darmstadt, Germany) using NdeI and EcoRI restriction enzymes, followed by ligase-based cloning. These vectors confer kanamycin (KANA) resistance and allow for high-level protein expression and purification in *E. coli.* LB medium, with or without 1% glucose and supplemented with KANA (50 µg/mL), was used as the standard growth medium. Plasmid maintenance and propagation were performed in *E. coli* DH5α cells (Stratagene, La Jolla, CA, USA), while protein expression was carried out in *E. coli* BL21 (DE3) strain (Agilent Technologies, Santa Clara, CA, USA). An empty pET42b(+) vector (referred to as the empty vector, EV) was generated by heat-shock transformation into *E. coli* and used as the negative control signal. Plasmids were purified using the GenElute™ Plasmid Miniprep Kit (Sigma-Aldrich, St. Louis, MO, USA) according to the manufacturer’s instructions. Plasmid sequences were verified by Sanger sequencing performed by Eurofins Genomics (Germany) using the TubeSeq Supreme protocol, and sequences were analysed and aligned using BioEdit (version 7.2.5).

### 2.3 KnowVolution of human HGD

#### 2.3.1 Generation of an HGD error-prone PCR library

Custom-designed primers containing 12 phosphorothioate (PTO) modifications at the 5’ end (Invitrogen, Thermo Fisher Scientific) were used for amplification of the HGD insert and the pET42b(+) plasmid backbone (Primer sequences in **Supplementary Data 2**). PCR of the vector backbone was performed in 50 µL reactions using Phusion High-Fidelity DNA polymerase supplemented with 1% DMSO, with the following cycling conditions: 98°C for 3 min; 35 cycles of 98°C for 30s, 58.9°C for 30s, and 72°C for 2.5 min; and a final elongation at 72°C for 5 min. Error-prone PCR (epPCR) of the HGD insert was carried out in 50 µL reactions using Taq DNA polymerase in the presence of MnCl_2_ (0.05-0.2 mM final concentration) to generate three independent mutant libraries, with cycling conditions of 94°C for 3 min; 35 cycles of 94°C for 30s, 53.3°C for 30 s, and 72°C for 1.5 min; and a final elongation at 72°C for 10 min. PCR products were analysed by agarose gel electrophoresis and digested with DpnI to remove methylated parental DNA, with vector backbones additionally treated with EcoRI to eliminate residual parental template DNA. Digested products were then column purified using the GenElute^TM^ PCR Clean-Up Kit (Sigma-Aldrich). For phosphorothioate-ligase independent cloning (PLICing) [29], purified vector and insert fragments were diluted to 0.03 and 0.09 pmol/µL (1:3 molar ratio) and subjected to iodine cleavage at 70 °C for 15 min using a fresh mixture of Tris-HCl, iodine in ethanol, and water (5:3:2 *v/v*). Cleaved fragments were hybridized at room temperature for 10 min, column purified and transformed into chemically competent *E. coli* BL21 (DE3) cells for expression, screening and selection, as described in Section 2.4.1.1. Religation and DpnI controls were included to monitor background colony formation. Variants of interest were plasmid-purified according to the manufacturer’s instructions and subjected to Sanger sequencing to identify the influential positions, and were subsequently re-screened using the workflow described in Section 2.4.1.2.

#### 2.3.2 Site-saturation mutagenesis library

Single-site saturation mutagenesis (SSM) was performed using the 22c-trick approach, which employs a mix of three primers to encode all 20 standard AAs while minimizing codon redundancy. The primer mix included NDT (12 AAs), VHG (9 AAs), and TGG (1 AA) primers at a 12:9:1 molar ratio, ensuring even AA representation and reducing the number of variants to be screened. An oversampling factor of three was applied to achieve ∼95% coverage of the theoretical variant space, requiring screening of ∼66 colonies per position [30–32]. SSM was carried out on eight beneficial positions identified from phase I of the KnowVolution approach (31, 54, 86, 91, 150, 223, 317, 354) and five additional catalytic site positions (336, 347, 359, 373, 401) based on the HGD mutation database [4]. Mutagenesis was performed on HGD-WT using a modified QuikChange™ protocol with an initial single-primer extension step to reduce primer dimer formation. Forward and reverse primer extension reactions were prepared separately, each consisting of the HGD-WT template, 22c-trick primer mix, Phusion High-Fidelity polymerase with 1% DMSO, and buffer. Primer sequences and position-specific annealing temperatures are listed in **Supplementary Table S2**. Reactions were cycled for three rounds (98 °C for 10 s, annealing at the position-specific temperature for 30 s, 72 °C for 60 s). Equal volumes of the forward and reverse primer reactions were then combined for a second PCR amplification, performed for 15 cycles (98 °C for 10 s, annealing at the same temperature for 30 s, 72 °C for 60 s) followed by a final extension at 72 °C for 10 min. PCR products were analysed by agarose gel electrophoresis, digested with DpnI to remove methylated parental DNA, and heat-inactivated. The resulting products, along with HGD-WT and EV controls, were transformed into chemically competent *E. coli* BL21 (DE3) cells for expression, screening, and selection, as described in Section 2.4.1.1. Variants showing improved activity were plasmid-purified and subjected to Sanger sequencing to identify the beneficial substitutions, and were subsequently re-screened using the workflow described in Section 2.4.1.2.

#### 2.3.3 Computer-assisted structural analysis

Five residues (Asn31, Ser54, Asp86, Gln354, Pro359) were selected for *in silico* SSM based on their predicted impact on HGD stability and catalytic activity. The CompassR strategy [33] guided the selection of substitutions compatible with improved enzyme performance by analysing the relative folding free energy change (ΔΔG_fold_) between mutant (mut) and WT enzymes, calculated as ΔΔG_fold_ = ΔG_fold,mut_ - ΔG_fold,WT_. Positive ΔΔG_fold_ values indicate decreased stability, and substitutions with ΔΔG_fold_ ≤ 0.36 kcal/mol were considered potentially beneficial. The crystal structure of HGD-WT (PDB ID: 1EYB, chain A, 1.9 Å resolution) was rotamerized and energy-minimized using YASARA’s “RepairObject” to correct non-standard torsion angles. ΔΔG_fold_ values were computed with FoldX (v5) in YASARA using five independent runs at 298 K, pH 7, and 0.05 M ionic strength. For each residue, PositionScan systematically substituted all 19 alternative AAs. Evolutionary conservation of HGD residues was analysed using ConSurf-DB [34]. Homologues were identified with HMMER [35] (E-value ≤ 0.0001), filtered to ≤ 95% identity with CD-HIT [36] (max 300 sequences, ≤ 10% overlap, ≥ 60% coverage), aligned with MAFFT [37], and a phylogenetic tree was constructed using Neighbor-Joining with maximum likelihood distances [38]. Site-specific evolutionary rates were calculated with Rate4Site [39], which accounts for phylogenetic relationships and provides credibility intervals. Conservation scores were projected onto the HGD sequence and visualized on the 3D structure using ChimeraX (v1.9) [40], enabling assessment of the structural and functional importance of the selected residues.

#### 2.3.4 Multi-site directed mutagenesis library

Combinatorial variants (see **Supplementary Table S3**) were generated from previously constructed single-point site-directed mutagenesis (SDM) variants using either Phusion Flash polymerase following the Q5 protocol (manufacturer’s instructions) or Phusion High-Fidelity polymerase, with or without DMSO (See **Table S3**). The PCR reaction mixture contained 0.4 µM primers (final concentration), and 0.4 ng/µL plasmid template DNA (final concentration). Primers were designed according to the method described by Liu and Naismith (see **Table S3**), using a design that promotes primer-template annealing and minimizes primer dimerization, thereby eliminating the need for the previously applied two-step QuikChange™ protocol [41]. PCR amplification was performed with an initial denaturation at 98 °C for 10 s, followed by 30 cycles of 98 °C for 1 s, annealing at position-specific temperatures (see **Table S3**) for 30 s, and extension at 72 °C for 4 min, followed by a final extension at 72 °C for 10 min. PCR products were analysed by agarose gel electrophoresis, digested with DpnI to remove methylated parental DNA, and subsequently heat-inactivated. Purified plasmids were verified by Sanger sequencing to confirm the intended substitutions prior to further use. Verified constructs were transformed into chemically competent *E. coli* BL21 (DE3) cells and screened using the workflow described in Section 2.4.1.2.

### 2.4 High-throughput screening assay

Unless otherwise specified, all experimental procedures were performed as previously described in Lequeue *et al.* [28]. The HGD activity assay was adapted to a 96-well plate format by miniaturizing protein expression from Erlenmeyer flasks to deep-well plates. Assay conditions were adjusted for HTS, with robustness reassessed after minor optimization. Detailed optimization procedures and robustness analyses are provided in **Supplementary Methods S1** and **S2**, respectively.

#### 2.4.1 HGD variant screening

##### 2.4.1.1 Primary library screening

Individual colonies of *E. coli* BL21 (DE3) carrying either HGD-WT (benchmark) or EV (negative control) plasmids, as well as individual colonies derived from epPCR libraries or SSM libraries, were inoculated into 300 µL of LB_KANA_ medium supplemented with 1% glucose in sterile 96-well deep-well plates (conical bottom) and incubated overnight at 30 °C with shaking at 330 rpm. Master glycerol stocks (25% *v/v*) were prepared for subsequent retrieval of selected variants. For protein expression, 75 µL of overnight culture was transferred into 525 µL of LB_KANA_ containing 1 mM Isopropyl β-D-1-thiogalactopyranoside (IPTG), followed by incubation for 24 h at 22 °C with shaking at 400 rpm. The induction temperature of 22 °C was selected based on previously optimized HGD expression conditions [42]. For high-throughput processing, 150 µL of each well of bacterial culture was transferred to a flat-bottom 96-well plate and centrifuged at 3200 x *g* for 10 min at 4 °C to pellet the cells. The supernatant was discarded, and the resulting cell pellets were frozen at −20 °C for at least 1 h. Cell lysis was performed by resuspending pellets in 80 µL of lysozyme solution (1 mg/mL in 50 mM Tris-HCl, pH 7.4) and incubating for 1 h at 37 °C with shaking at 400 rpm in a Thermomixer (Eppendorf). Following lysis, clarified supernatants were transferred to a separate 96-well plate preloaded with assay mixture consisting of 50 mM potassium phosphate buffer (pH 7.4), 2 mM ascorbate, 0.2 mM Fe_2_SO_4_, and 2.5 mM HGA as substrate. Enzymatic activity was measured using an enzyme-to-assay mix ratio of 25:75 (*v/v*) in a final volume of 100 µL. This ratio showed the highest linearity compared with the other conditions tested (**Fig. S1a & S1b**). Each variant was assayed in a single measurement, while HGD-WT and EV controls were included in four technical replicates each. HGD activity was determined by monitoring the conversion of HGA to MAA at 340 nm, with absorbance recorded every 30 s for 25 min at 37 °C, and enzyme velocities and residual activities were calculated exactly as described in our previously published method [28]. Reaction linearity was confirmed for all variants and controls prior to calculation. Background signals were normalized to the empty vector (EV) negative control [28].

##### 2.4.1.2 Confirmation re-screening of selected variants

Variants exhibiting improved catalytic activity in the initial high-throughput screen were selected for activity confirmation, and recombinatorial variants generated by SDM during Phase IV were likewise evaluated using the re-screen protocol. Selected clones, along with HGD-WT and EV controls, were inoculated from the frozen master plate using sterile toothpicks or, in the case of combinatorial variants, from glycerol stocks. Each variant was grown overnight in LB_KANA_ supplemented with 1% glucose at 37 °C with shaking at 250 rpm. The following day, cultures were diluted into fresh LB_KANA_ to an initial OD_600_ of 0.1, measured using a spectrophotometer, and incubated at 30 °C with shaking at 250 rpm until mid-log phase (OD_600_ 0.6-0.8) was reached. Protein expression was induced with 1 mM IPTG. After induction, 600 µL of each culture was transferred into 96-well deep-well plates (conical bottom) in eight technical replicates per variant, HGD-WT, or EV, and incubated for 24 h at 22 °C with shaking at 400 rpm. Following incubation, 150 µL of each culture was collected for cell lysis. Enzymatic activity was measured using an enzyme-to-assay mix ratio of 10:90 (*v/v*), which provided optimal linearity under the slightly different protein expression conditions of the re-screen (**Fig. S1b**). Absorbance was monitored every 30 s for 20 min at 37 °C. Reaction linearity was confirmed for all variants prior to calculation, and enzyme velocities and relative activities were calculated according to our previously published method [28].

##### 2.4.1.3 Variant selection and activity thresholds

For Phase I of KnowVolution, the proportion of active clones (in %) was calculated to assess library quality and optimize the balance between mutation load and retention of enzymatic function, ensuring that the libraries contained a sufficient number of functional variants for effective discovery of influential positions. Variants were classified as active if their enzymatic activity exceeded the mean activity of the EV plus three standard deviations (SDs) (Mean + 3xSD), and the percentage of active clones was calculated as the number of active variants divided by the total number of variants, multiplied by 100. Candidate variants were initially identified in the primary HTS based on apparent activity exceeding that of the HGD-WT enzyme, thereby highlighting AA positions with potential functional impact. These candidates were subsequently evaluated in a more accurate re-screen assay to confirm activity. Variants retaining at least 85% of WT activity in the re-screen were selected for further analysis. This threshold was chosen to capture positions that, while not exceeding HGD-WT activity, might still influence catalytic function and could be optimized in subsequent phases. This approach ensures that both clearly beneficial and subtly influential positions are included, providing a solid basis for targeted saturation mutagenesis and combinatorial optimization in the KnowVolution workflow. For Phases II and IV of KnowVolution, only variants with reproducibly higher mean activity than HGD-WT (> 100%) were selected for further analysis.

##### 2.4.1.4 Statistical analysis

Outliers were detected and excluded using the ROUT test. Data normality was evaluated with the Shapiro-Wilk test. When the dataset followed a normal distribution, differences in % residual activity (calculated from enzyme velocity) between HGD variants and the HGD-WT control were analysed using a one-way parametric ANOVA, followed by Bonferroni post-hoc correction to account for multiple comparisons. For datasets that were not normally distributed, the Kruskal-Wallis test followed by Dunn’s post-hoc test was applied to identify statistically significant differences between each variant and the HGD-WT control. All statistical analyses were performed in GraphPad Prism version 10.3.1.

### 2.5 Structural modelling and visualization

The hexameric structure of human HGD and the docking of the catalytic cofactor Fe^2+^ and substrate HGA in chain A were predicted using the Boltz2 [43] server based on the human HGD AA sequence (UniProt ID: Q93099). The resulting protein-ligand complex was visualized and further analysed in UCSF ChimeraX (version 1.9) [40]. Identified residues from phase I of KnowVolution were classified into three categories based on their spatial position. Catalytic pocket residues were defined as residues with atoms within 5 Å of either Fe^2+^ or HGA. Surface residues were defined as solvent-exposed residues, determined by visual inspection after displaying the protein surface. All remaining buried residues, neither solvent-exposed nor within 5 Å of the ligands, were classified as core residues. Interface analysis of the HGD biological assembly was performed using PDBePISA (v1.52, [20/10/2014]). Three key parameters were considered: solvent-accessible surface area (ASA), buried surface area (BSA), and interface ΔG. Only interfaces with a complex significance score (CSS) of 1 were considered biologically relevant [44–47]. Following Phase IV, selected HGD variants were analysed to investigate structural factors that might explain differences in enzymatic activity compared to HGD-WT. Multiple-residue substitutions corresponding to the selected experimentally tested Phase IV recombinants were introduced in UCSF ChimeraX (version 1.9) to generate structural models of the variant enzymes. Tunnel and access pathway analyses were performed using CAVER Web [48] to assess whether the observed activity differences could be associated with alterations in substrate or product channels. Representative high-activity and low-activity recombinants were selected for comparison with HGD-WT. This approach allowed evaluation of potential differences in tunnel geometry, accessibility, and connectivity that could contribute to the functional effects of the substitutions. Complete CAVER Web outputs are provided in the Data Availability section. In addition, the putative active-site pockets of HGD-WT and selected high-activity and low-activity recombinants were analysed using CASTpFold [49]. The same ChimeraX-generated structural models, including the hexameric HGD complex with Fe²⁺ and HGA in chain B, were used as input. For each structure, CASTpFold calculated pocket volume, surface area, and the residues lining the pocket. Pockets at the hexameric and dimeric interfaces as well as the catalytic pockets of each protomer were selected for comparative analysis across variants. Complete CASTpFold outputs, including full residue lists for each pocket, are provided in the Data Availability section. In addition, putative flexible loop regions were analysed using LoopGrafter [50] to support structural interpretation of mutation sites and their surrounding regions. HGD-WT was used as the scaffold structure, while the P359E+Q354P recombinant was used as the insert structure.

## 3 Results and discussion

AKU arises from HGD deficiency, and although NTBC offers palliative metabolic control, it does not restore HGD enzymatic activity, thereby motivating further investigation into HGD structure-function relationships for the development of restorative therapies [9–11]. To address this, KnowVolution was employed as a controlled approach to identify substitutions that improve enzyme stability and catalytic efficiency, while enabling detailed insights into structure-function relationships with potential added value for precision medicine approaches in AKU. The strategy combines random and targeted mutagenesis with computational analysis and rational recombination to efficiently explore the protein sequence space, uncover cooperative effects between residues, and gain mechanistic insights that traditional DPE often misses (**Fig. 2**) [27].

**Figure 2.**
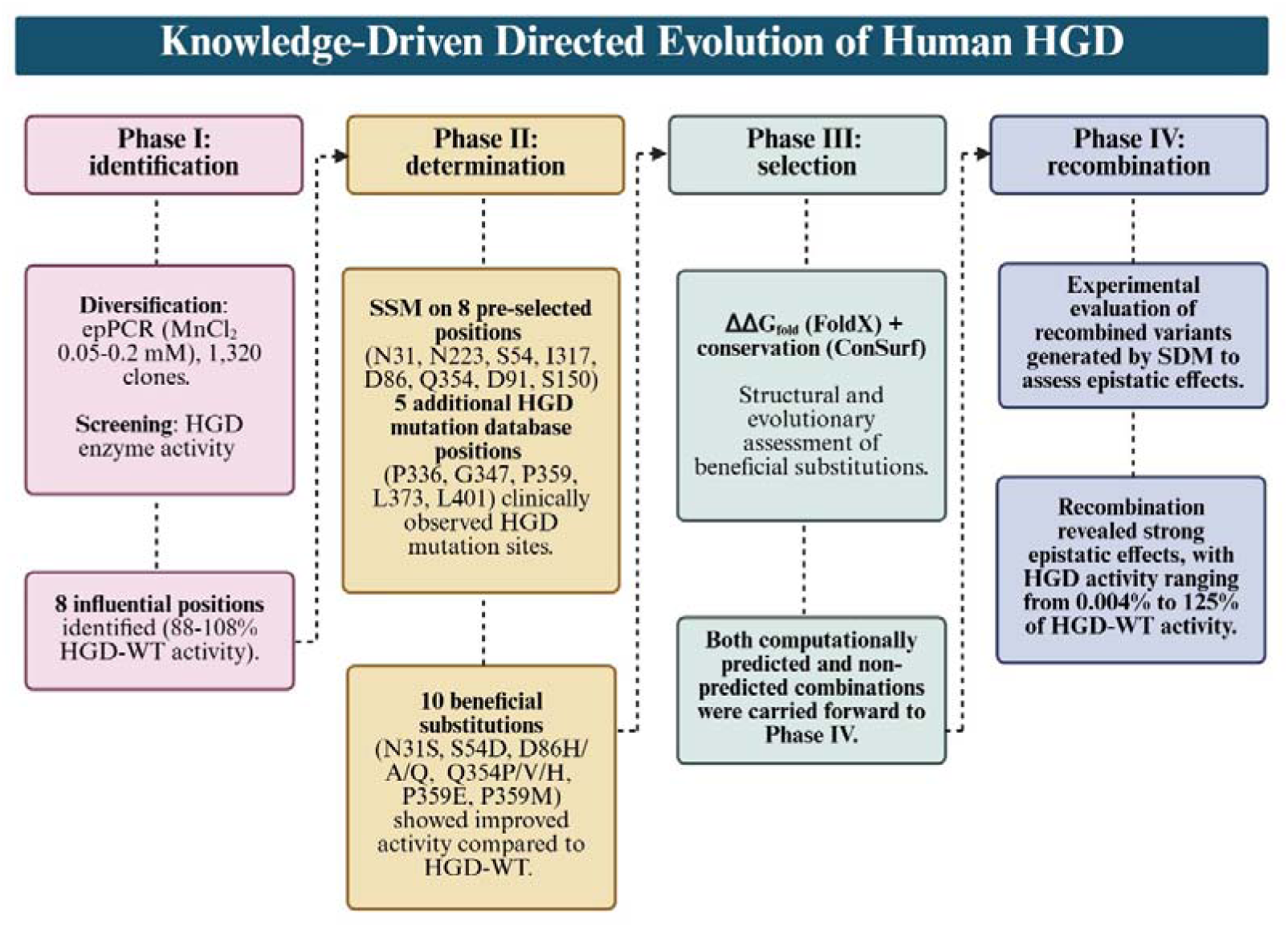
Knowledge gaining directed evolution of human HGD. Phase. **I**: Diversification of HGD by epPCR generated three libraries with increasing mutational load (MnCl_2_ concentrations: 0.05, 0.1, 0.2 mM), which were screened to identify eight positions with potentially influential effects (88-108% relative activity vs. HGD-WT). **Phase II**: SSM was performed at the eight Phase I positions plus five additional positions from the HGD mutation database, yielding ten substitutions that improved catalytic activity (N31S, S54D, D86H/A/Q, Q354P/V/H, P359E/M). **Phase III**: Computational analysis using FoldX (ΔΔG_fold_) and ConSurf scores assessed the energetic and evolutionary constraints of these substitutions to guide recombination in Phase IV. **Phase IV**: Combinatorial variants were generated from the most promising Phase II substitutions, selected based on structural proximity, favourable ΔΔG_fold_ (≤ 0.36 kcal/mol), and evolutionary conservation. A total of 22 recombinants were constructed *via* SDM and screened to experimentally evaluate epistatic effects, revealing strong interactions with activities ranging from 0.004% to 125% relative to HGD-WT.

To ensure reliable assessment of variant activity, the high-throughput HGD activity assay was optimized and evaluated for robustness prior to library screening. Details of assay optimization and assay robustness, including plate uniformity and performance metrics, are presented in **Supplementary Results S1** and **S2**, respectively, and summarized in **Table S1**.

In Phase I of KnowVolution, the focus was on identifying AA positions that contribute to catalytic performance while preserving overall enzyme function, with variants retaining at least 85% of HGD-WT activity selected as starting points for subsequent optimization. In Phase I of the workflow, random mutagenesis was performed by epPCR to generate three libraries with increasing MnCl_2_ concentrations (0.05, 0.1, and 0.2 mM), yielding 440, 528, and 352 clones, respectively, for a total of 1,320 variants. Under these conditions, the proportion of active clones decreased with higher mutagenesis: 92.3 ± 2.2%, 80.1 ± 11.6%, and 71.6 ± 14.8% remained active in the 0.05, 0.1, and 0.2 mM libraries, respectively, indicating that higher mutation rates reduced functional variants while increasing sequence diversity. In the primary screen, 24 clones from the 0.05 mM library, six from the 0.1 mM library, and eight from the 0.2 mM library exhibited activity at or above the threshold of 85% relative to HGD-WT. Re-screening of active variants yielded 16 clones representing eight unique AA positions. Their relative activities (mean ± SD) ranged from 88.1 ± 9.4% for N31S to 108.3 ± 4.5% for N223D/S54N, with additional variants such as I317V (104.5 ± 10.3%), D86G (104.9 ± 2.4%), Q354R (95.5 ± 16.6%), D91G (102.2 ± 3.0%), and S150I (99.7 ± 3.4%) showing activities between these extremes (**Fig. 3a**). To gain structural and mechanistic insight, the identified positions were mapped onto the 3D monomeric structure of HGD. Four residues were located within the protein core (Asn54, Ser150, Asn223, and Ile317) and four on the surface (Gln354, Asp86, Asp91, and Asn31). Notably, none of the identified positions were located within the catalytic pocket (**Fig. 3b**).

**Figure 3.**
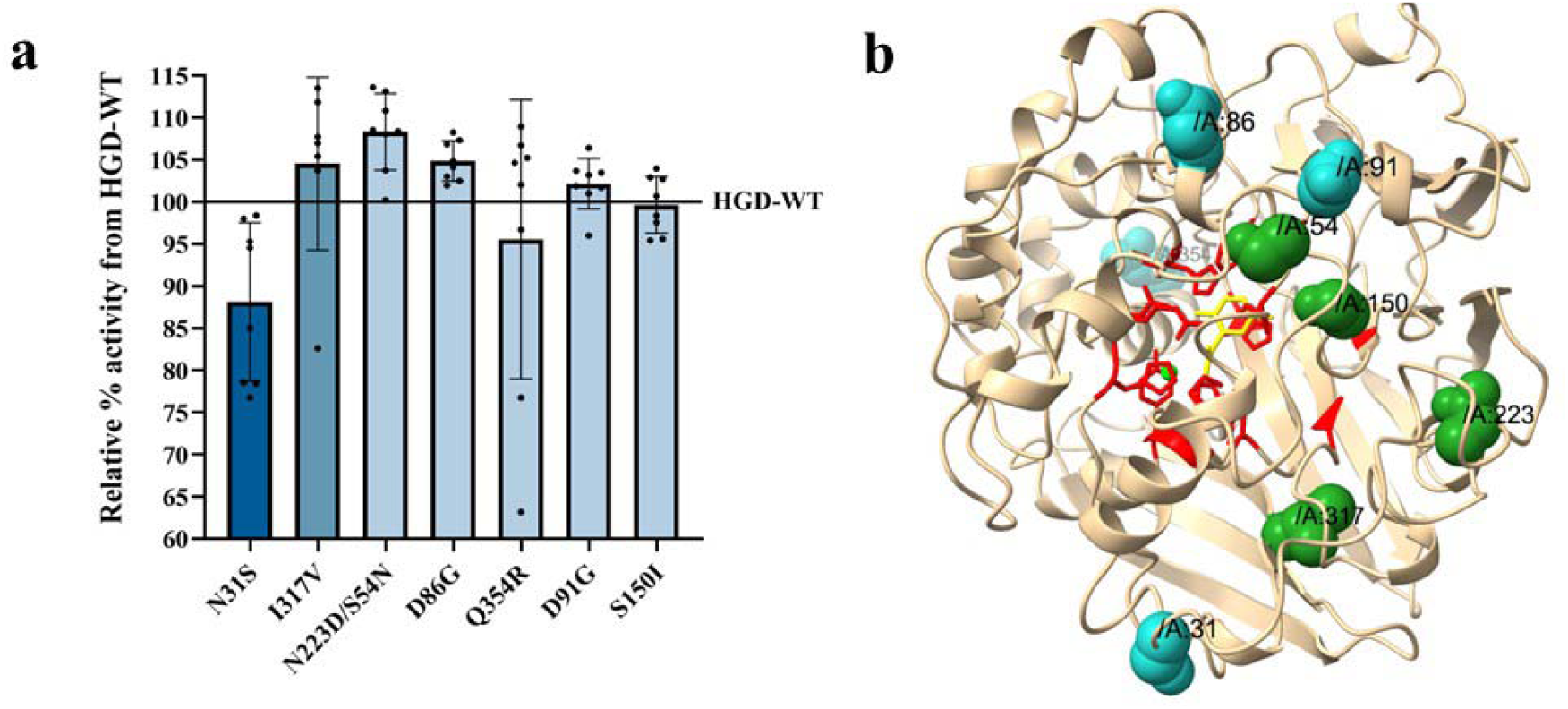
Identification and structural mapping of influential HGD positions from Phase I of KnowVolution. **(a)** epPCR libraries were generated with increasing MnCl□ concentrations (0.05, 0.1, 0.2□mM; visualized as a blue gradient from light to dark) to introduce variants with progressively higher mutational load. Variants retaining ≥□85□% of HGD-WT activity were selected, identifying positions with potential to enhance catalysis in subsequent phases of the KnowVolution workflow. Eight unique positions were chosen for Phase II SSM, with relative activities ranging from 88.1□±□9.4□% (N31S) to 108.3□±□4.5□% (N223D/S54N) of WT. Relative activities are expressed as mean ± SD compared to HGD-WT (100%). Outliers were excluded using the ROUT test. **(b)** Structural mapping of residues identified in Phase I on chain A of the human HGD monomer. The catalytic pocket (defined as residues located within 5 Å of both the cofactor and substrate) is shown in red, residues in the protein core in green, and surface residues in blue. Core positions included Asn54, Ser150, Asn223, and Ile317, while surface positions included Gln354, Asp86, Asp91, and Asn31. The substrate (HGA) is shown in yellow, and the non-heme Fe^2+^ cofactor is shown in fluorescent green. None of the residues were located within the catalytic pocket.

Building on the influential positions identified in Phase I, Phase II systematically evaluated AA substitutions at sites likely to influence HGD activity. The eight positions from Phase I (31, 54, 86, 91, 150, 223, 317, and 354) were targeted using SSM, along with five additional positions (336, 347, 359, 373, and 401) from the HGD mutation database [4]. Inclusion of these patient-derived hotspots allowed us to obtain experimental evolution data at sites known to impact AKU, enabling the exploration of alternative substitutions that could enhance or modulate catalytic function. In total, 13 SSM libraries were generated using the 22c-trick strategy, and 66 clones per library were screened to achieve ∼95% theoretical coverage. Variants showing improved activity (≥ 100% of HGD-WT) were re-screened to confirm reproducibility. Of the 13 targeted positions, substitutions at five sites (Asn31, Ser54, Asp86, Gln354, and Pro359) produced variants with increased activity relative to HGD-WT. At position Asp86, three beneficial substitutions were identified (D86H, D86A, D86Q), while three substitutions were obtained for Gln354 (Q354P, Q354V, Q354H), and two for Pro359 (P359E, P359M). The corresponding mean relative activities (mean ± SD) were: N31S, 105.9 ± 9.0%; S54D, 111.9 ± 13.9%; D86H, 107.0 ± 2.8%; D86A, 103.3 ± 6.3%; D86Q, 102.1 ± 8.7%; Q354P, 100.2 ± 7.9%; Q354V, 108.0 ± 10.2%; Q354H, 113.7 ± 5.4%; P359E, 119.9 ± 9.5%; and P359M, 106.0 ± 12.2%. Among these, four variants (S54D, D86H, Q354H, and P359E) showed statistically significant increases in activity compared to HGD-WT (S54D, p = 0.0079; D86H, p = 0.0423; Q354H, p = 0.0109; P359E, p = 0.0001) (**Fig. 4a**).

**Figure 4.**
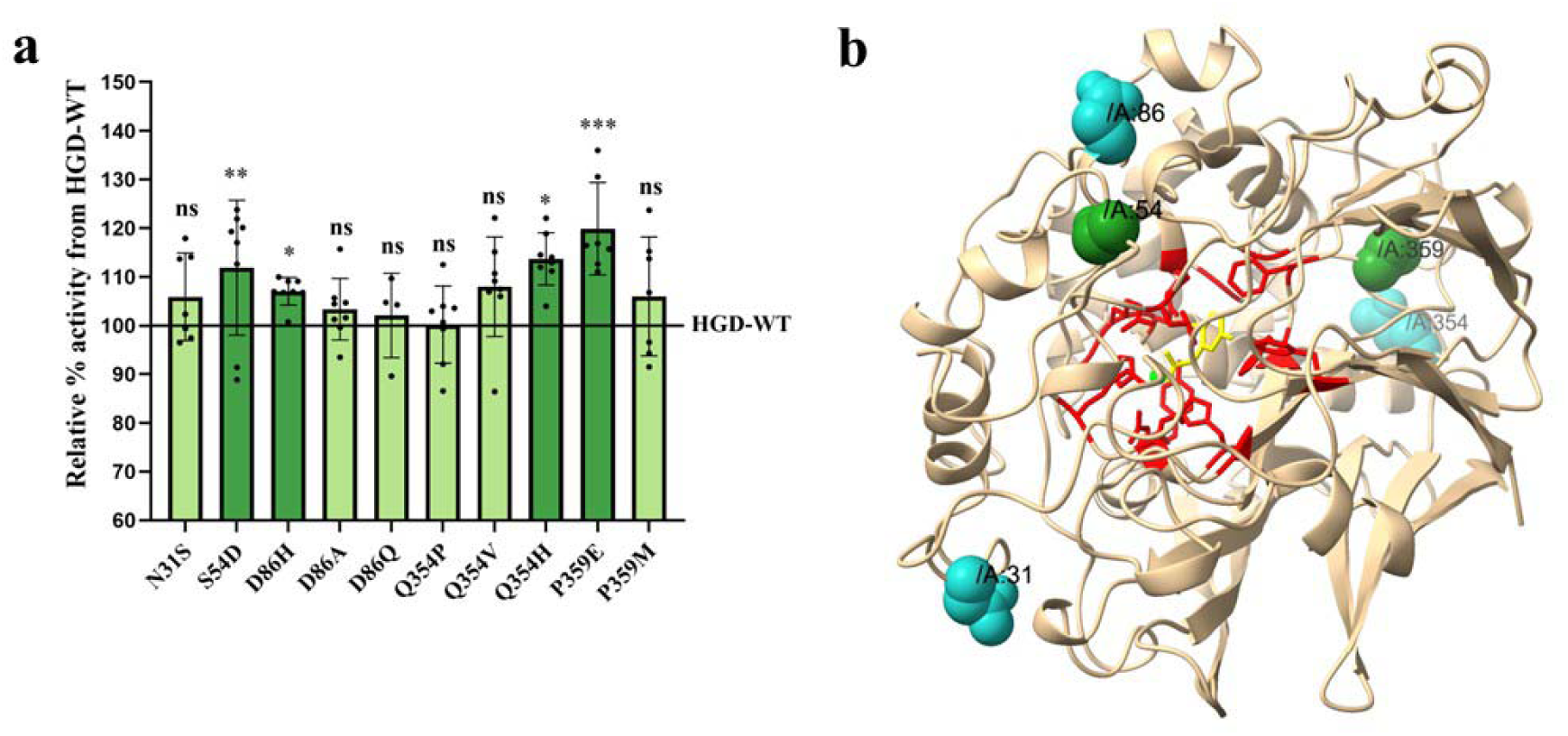
Beneficial substitutions from Phase II screening and their structural mapping within the monomeric 3D structure of human HGD. **(a)** Thirteen SSM libraries were generated at the eight positions identified in Phase I, along with five additional positions from the HGD mutation database. Variants were screened, and mean relative activities ranged from ∼100□% to ∼120□% of HGD-WT (100□%), visualized as a green gradient from light green (lower activity) to dark green (higher activity). Relative activities are expressed as mean□±□SD compared to HGD-WT. Outliers were excluded using the ROUT test. Statistical analysis was performed using one-way ANOVA with Bonferroni post-hoc correction or Kruskal-Wallis with Dunn’s post-hoc test depending on normality. Significance levels: p□≤□0.05 (*), p ≤ 0.01 (**), p□≤□0.001 (***). **(b)** Structural mapping of residues showing beneficial substitutions in Phase II on chain A of the human HGD monomer, using the same color scheme: catalytic pocket (red), core (green), and surface (blue), substrate (yellow), cofactor (fluorescent green). Beneficial substitutions were identified at Asn31 (surface), Ser54 (core), Asp86 (surface), Gln354 (surface), and Pro359 (core). None of these residues were located within the catalytic pocket.

To place these activity improvements in a structural context, the identified positions were mapped onto the 3D monomeric structure of HGD (**Fig. 4b**). Notably, the beneficial substitutions were distributed across structurally distinct regions of HGD: Asn31 is a solvent-exposed residue on the N-terminal loop, Asp86 is surface-exposed within the flexible 77-114 loop of the N-terminal domain, Ser54 is moderately solvent-accessible within the N-terminal β-sheet forming part of the structural β-sandwich, Gln354 is surface-exposed in the C-terminal β-sheet, and Pro359 is buried in the core of the C-terminal β-sheet [12,40]. Importantly, none of the identified positions are located within the catalytic pocket, suggesting that their effects on enzymatic activity are indirect rather than due to direct participation in catalysis. Several of these positions are solvent-exposed, supporting the notion that their functional impact is more likely related to structural stabilization or subunit interactions. Human HGD assembles into a hexameric complex stabilized by intersubunit β-sheet contacts, hydrophobic interactions, and defined loop positioning. Crystal structures further indicate that this hexameric assembly is representative of the solution state, as lattice contacts are predominantly water-mediated and the solvent content is high (∼68%). Proper hexamer formation is therefore essential for enzymatic function, since accurate subunit orientation and loop positioning are required to maintain the geometry of the active site. Accordingly, several AKU-causing mutations occur at surface or interface positions and destabilize the oligomeric assembly [4,12]. To further assess the involvement of the identified positions in subunit interactions, interface analysis of the HGD biological assembly was performed using PDBePISA [47,51,52]. Among the five positions yielding beneficial substitutions, Asn31 and Asp86 showed measurable contributions to the interface. Asn31 exhibited a solvent-accessible surface area (ASA = 95.70 Å²), a buried surface area upon complex formation (BSA = 89.87 Å²), and a favourable free energy contribution (ΔG = −0.24 kcal/mol), consistent with a stabilizing role at the subunit interface. Asp86 showed high solvent exposure (ASA = 139.29 Å²) and moderate burial (BSA = 40.81 Å²), with a stabilizing contribution (ΔG = −0.19 kcal/mol). Structurally, Asp86 forms one hydrogen bond and two salt bridges with Lys126, involving the OD1 and OD2 atoms of Asp86 and the NZ atom of Lys126 at distances of 2.88 and 3.68 Å, respectively, highlighting its potential role in stabilizing the oligomer. Although Ser54 displayed substantial solvent exposure (ASA = 95.08 Å²) without detectable BSA or ΔG contributions in PDBePISA analysis, structural inspection in ChimeraX[40] revealed direct interactions between symmetry-related Ser54 residues across adjacent chains at the dimeric interface. This observation suggests that Ser54 may still contribute to oligomer stabilization through intersubunit polar contacts, that are not sufficiently captured by energetic interface calculations. In contrast, Pro359 exhibited limited solvent accessibility (ASA = 42.87 Å²) without measurable BSA or ΔG contributions, and no interface parameters were obtained for Gln354, indicating the absence of detectable interface contacts at these positions. In contrast, the beneficial effects observed for Gln354 and Pro359 are more likely related to indirect effects on protein stability or local structural dynamics rather than direct interface interactions or catalytic changes.

Taken together, these results indicate that the beneficial substitutions identified by the KnowVolution campaign predominantly target surface and structural regions outside the catalytic pocket. While Asn31 and Asp86 show clear energetic contributions to the subunit interface, structural analysis additionally suggests a possible role for Ser54 in dimeric interface stabilization. The remaining substitutions likely enhance catalytic performance through indirect, non-catalytic mechanisms involving structural stabilization and altered protein dynamics. Similar effects of distal substitutions on catalytic efficiency have been reported for other enzyme systems [53–55].

During Phase III, we assessed the energetic and evolutionary constraints of the beneficial substitutions by predicting single-residue ΔΔG_fold_ values with FoldX (CompassR) and analyzing residue conservation using ConSurf, both performed during Phase III to guide Phase IV recombination experiments [33,34]. Predicted ΔΔG_fold_ values were: N31S (0.26), S54D (−0.06), D86A (0.04), D86Q (0.02), D86H (0.23), Q354V (0.02), Q354P (0.16), Q354H (0.24), P359M (2.03), and P359E (0.90) (**Fig. 5a**). ConSurf scores indicated high conservation for Asn31 and Pro359 (7-8), low for Ser54 and Asp86 (1), and moderate for Gln354 (5), highlighting differential evolutionary constraints. The 3D structure of chain A of HGD-WT is shown in **Fig. 5b** with the ConSurf colour scale overlaid, where bluer colours indicate more variable positions and pinker colours indicate more conserved positions. This visualization indicates that Asn31 and Pro359 are highly conserved, whereas Ser54 and Asp86 are more variable. Chemically, substitutions span conservative, semi-conservative, and non-conservative changes: N31S and D86Q are conservative polar-to-polar, D86H and Q354H are semi-conservative polar/charged substitutions, and S54D, D86A, Q354P, and P359E represent non-conservative changes altering charge, polarity, or rigidity. FoldX predictions indicated that all substitutions are energetically tolerated (ΔΔG_fold_ ≤ 0.36 kcal/mol), even at conserved positions such as Asn31 and Pro359, consistent with the observed activity gains and suitable for Phase IV recombination. For instance, the non-conservative P359E substitution at a highly conserved buried position increased activity by ∼20%, demonstrating that even constrained sites can tolerate beneficial changes if compatible with the fold and not directly involved in catalysis. Alternatively, residues that are less evolutionarily conserved or located on the protein surface, such as Ser54, Asp86, and Gln354, were able to accommodate a variety of substitutions without significantly affecting protein stability, allowing these changes to enhance activity without disrupting the overall fold.

**Figure 5.**
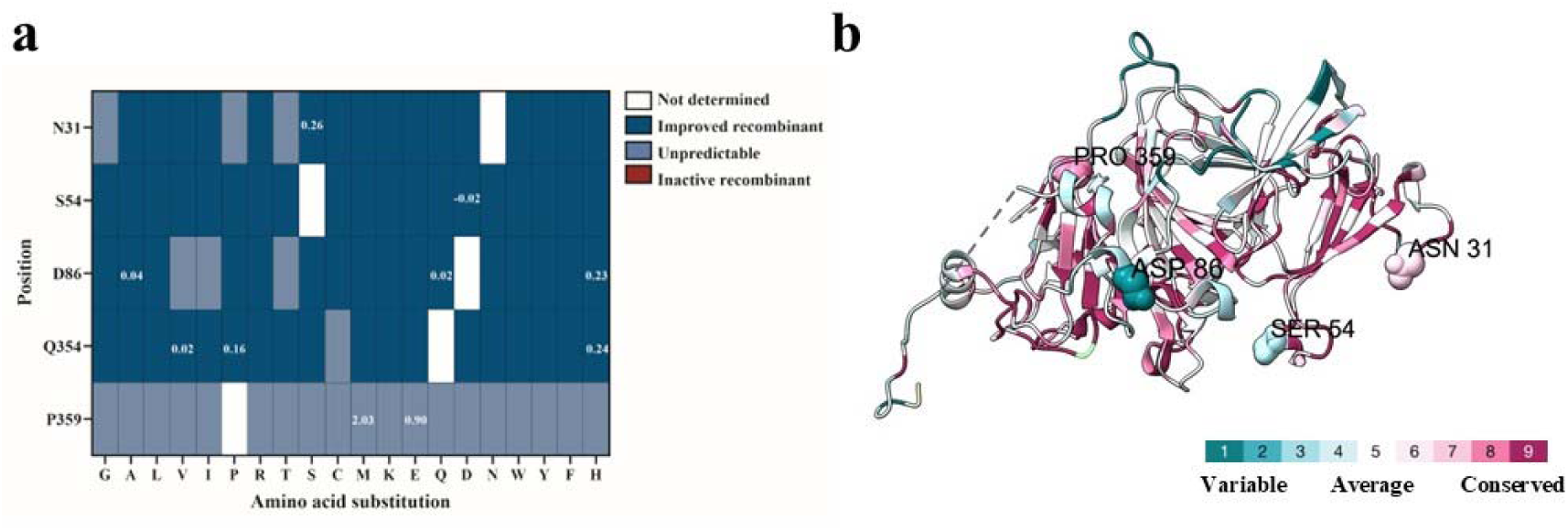
Phase III evaluation of beneficial HGD substitutions identified in Phase II. **(a)** Single-residue stability predictions were performed using the FoldX algorithm within the CompassR approach. All possible substitutions at the five beneficial positions (Asn31, Ser54, Asp86, Gln354, and Pro359) were evaluated by calculating ΔΔG_fold_ values. Substitutions were classified as energetically compatible (ΔΔG_fold_ ≤ 0.36 kcal/mol; dark blue), uncertain (0.36-7.52 kcal/mol; light blue), or predicted to be inactivating (≥ 7.52 kcal/mol; red). The experimentally identified beneficial substitutions (N31S, S54D, D86H/A/Q, Q354P/V/H, and P359E/M) are indicated in their corresponding boxes and all fall within the energetically compatible range, supporting their suitability for recombination in Phase IV. **(b)** Structural mapping of evolutionary conservation scores obtained from ConSurf-DB onto chain A of the HGD-WT structure, visualized in ChimeraX. Residues are coloured from blue (variable; score 1) to pink (highly conserved; score 9). Beneficial positions are highlighted, showing that Asn31 and Pro359 are highly conserved, whereas Ser54 and Asp86 are more variable. Gln354 could not be visualized due to its absence in the ConSurf-derived structural model.

In Phase IV, we combined substitutions that had shown the most promising activity improvements in Phase II, selecting combinations based on structural proximity, predicted energetic compatibility (ΔΔG_fold_ ≤ 0.36 kcal/mol), and evolutionary conservation. A total of 22 recombinant variants were constructed *via* SDM and subsequently screened. Their corresponding mean relative activities (mean ± SD) were as follows: N31S+S54D (115.9 ± 9.3%), N31S+S54D+D86Q (111.7 ± 9.4%), N31S+S54D+D86A (118.6 ± 17.2%), N31S+S54D+D86H (125.0 ± 13.7%), S54D+D86Q (101.2 ± 6.0%), S54D+D86H (0.004 ± 0.01%), S54D+D86A (101.4 ± 8.9%), S54D+Q354P (100.2 ± 14.7%), S54D+Q354V (96.9 ± 10.3%), S54D+Q354H (105.3 ± 11.6%), D86Q+Q354P (70.3 ± 5.6%), D86Q+Q354V (76.2 ± 9.2%), D86Q+Q354H (0.25 ± 0.36%), D86H+Q354P (101.5 ± 2.3%), D86H+Q354V (104.8 ± 5.3%), D86H+Q354H (118.2 ± 13.9%), P359E+Q354P (111.1 ± 8.6%), P359M+Q354P (104.8 ± 8.0%), P359E+Q354H (99.7 ± 4.0%), P359M+Q354H (96.0 ± 6.2%), P359E+Q354V (73.7 ± 3.8%), and P359M+Q354V (93.9 ± 8.3%) (**Fig. 6a**). Among these, ten recombinants exhibited statistically significant differences in activity compared with HGD-WT, including increased activity (N31S+S54D, p = 0.0048; N31S+S54D+D86A, p = 0.0010; N31S+S54D+D86H, p < 0.0001; D86H+Q354H, p = 0.0008; P359E+Q354P, p = 0.0055) and decreased activity (S54D+D86H, p < 0.0001; D86Q+Q354P, p < 0.0001; D86Q+Q354V, p < 0.0001; D86Q+Q354H, p < 0.0001; P359E+Q354V, p < 0.0001). Because none of the recombined positions are located within the catalytic pocket, we next investigated whether the observed differences in catalytic activity could be related to substrate or product access. To this end, tunnel analysis was performed using CAVER Web on selected high- and low-activity recombinants, with HGD-WT serving as a reference [48]. The analysis revealed that the overall architecture of the main access pathway (*i.e.*, the primary tunnel identified by CAVER that connects the protein surface to the buried active site and is likely to facilitate substrate entry and product release) remains largely unchanged across all variants. The bottleneck radius was identical (1.8 Å) in both high- and low-activity recombinants, matching the WT enzyme, while tunnel lengths varied only slightly (17.1-19.4 Å). Similarly, throughput values remained essentially constant, and the set of residues lining the tunnel was unchanged. These observations suggest that diffusion through the primary tunnel is unlikely to drive the observed activity differences. Instead, these results suggest that the effects of the mutations arise from local structural or dynamic changes rather than alterations in the main access pathway. For reference, the tunnel architecture of HGD-WT, shown in complex with ligand and cofactor, is illustrated in **Fig. 6b**, while quantitative tunnel parameters are summarized in **Table S4**. To further evaluate potential structural contributions to activity, pockets at the hexameric and dimeric interfaces as well as the catalytic pockets of each protomer were analysed using CASTpFold [49]. Some recombined positions, notably residues 54 and 86, form part of the pocket enclosed by the hexameric interface (see Data Availability for CASTpFold job IDs and complete residue lists). For reference, the putative hexameric interface, dimeric interface and active-site pockets of HGD-WT are illustrated in **Fig. 6c,d**, while quantitative pocket parameters are shown in **Fig.6e-j**. High-activity variants containing either S54D or D86H showed reduced solvent-accessible pocket volume and surface area at the hexameric interface compared to HGD-WT (**Fig. 6e,f**). Pocket volumes decreased by 4.7-6.6%, and pocket surface areas decreased by approximately 1.5%. However, no consistent relationship between these changes and catalytic activity was observed, as the double mutant S54D+D86H displayed low activity (**Fig. 6a**). Substitution of His86 with Gln in a high-activity variant containing Q354H restored hexameric interface pocket volume and surface area to WT-like values and was associated with low catalytic activity (**Fig. 6a,e,f**). No changes in volume or surface area were detected for pockets at the dimer interface across all analysed variants (**Fig. 6g,h**). In contrast, one high-activity variant (Q354P+P359E) exhibited reduced surface area and volume across all six catalytic pockets compared to HGD-WT (**Fig. 6i,j**). Interestingly, residues 354 and 359 are located in close spatial proximity to the Gly347-Gly355 loop previously described in the literature [14], which acts as a dynamic lid over the active-site pocket and shields it from solvent upon HGA binding, underscoring the potential relevance of this region to the catalytic effects observed here. In line with this, LoopGrafter [50] analysis identified a putative loop region spanning residues 340-365, encompassing both residues 354 and 359, and revealed only minor geometric differences between HGD-WT and the Q354P+P359E variant. The overall loop geometry remained largely conserved, as reflected by the nearly identical end-to-end distance parameter (D = 11.89 in HGD-WT versus 11.92 in the variant). In contrast, small increases in the δ and θ orientation parameters (133.89 to 135.7 and 136.08 to 138.76, respectively), together with a decrease in the ϱ packing parameter (29.08 to 26.7), suggest subtle changes in loop orientation and local packing rather than major conformational rearrangements. These observations indicate that local dynamics within this region may contribute to the functional effects associated with these substitutions. These predictions could be further refined and complemented by additional computational approaches, such as molecular dynamics simulations or electrostatic surface analyses, which would also help to address the inherent limitations of static structure-based tools, particularly in capturing highly flexible loop regions associated with active-site function.

**Figure 6.**
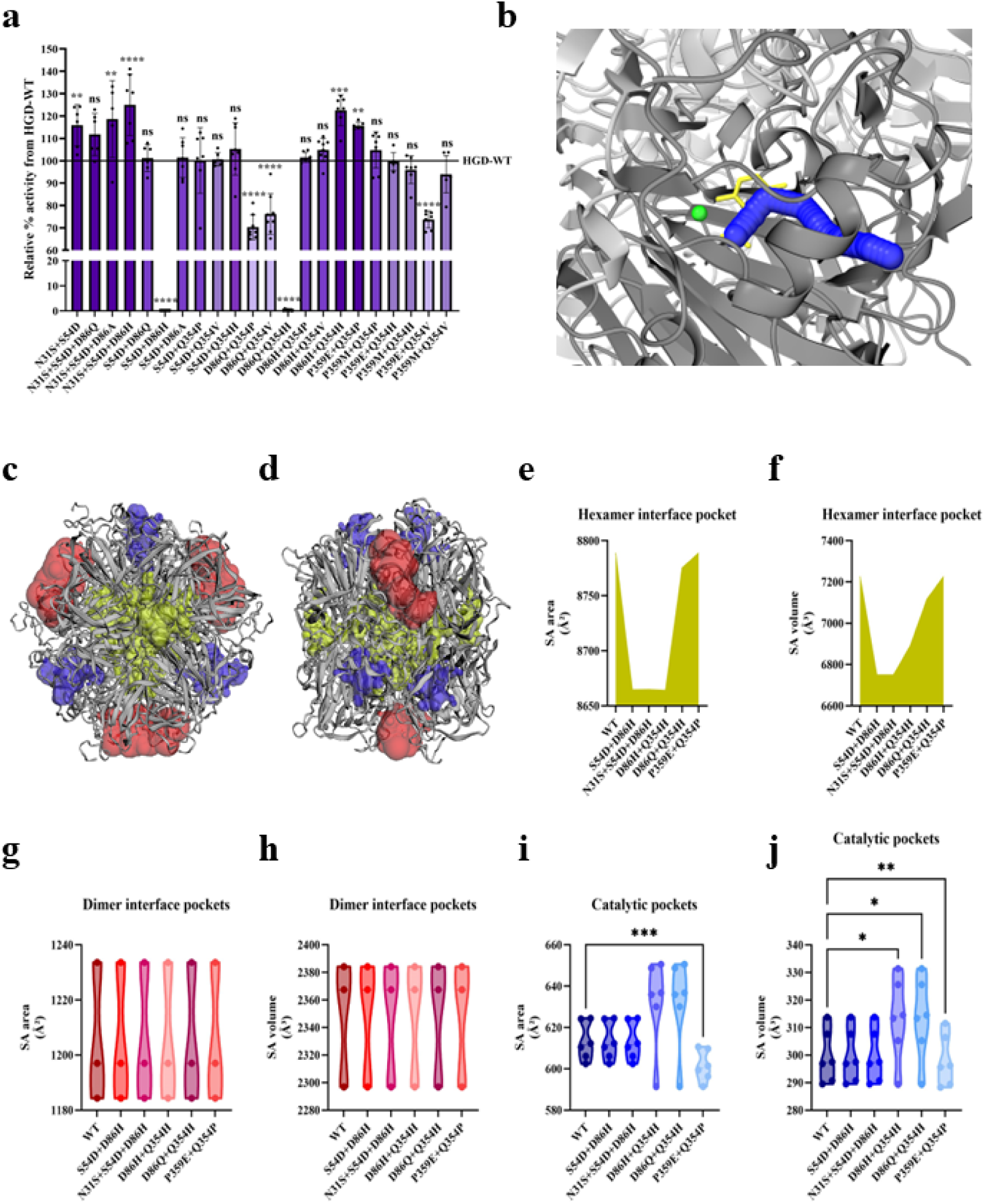
Structural and functional analysis of Phase IV recombinants. **(a)** Phase IV recombination of selected substitutions guided by structural proximity, predicted energetic compatibility (ΔΔG_fold_ ≤ 0.36 kcal/mol), and evolutionary conservation. A total of 22 recombinant variants were generated by SDM and experimentally screened. Mean relative activities ranged from ∼0.004 % to ∼125 % of HGD-WT (100 %) and are visualized using a purple gradient from light (low activity) to dark (high activity). Activities are reported as mean ± SD. Outliers were excluded using the ROUT test. Statistical significance was assessed using one-way ANOVA with Bonferroni post hoc correction or Kruskal–Wallis with Dunn’s post hoc test, depending on data normality (**p ≤ 0.01, ***p ≤ 0.001, ****p ≤ 0.0001). **(b)** Tunnel analysis performed with CAVER Web on selected high- and low-activity recombinants, using HGD-WT as a reference. The main access pathway shows highly conserved geometry across all variants, with identical bottleneck radii (1.8 Å), minimal variation in tunnel length (17.1-19.4 Å), unchanged throughput values, and a conserved set of tunnel-lining residues. These results indicate that diffusion through the primary tunnel is unlikely to account for the observed activity differences. For reference, the tunnel architecture of HGD-WT (chain B) is shown in blue, with the ligand HGA in yellow and the Fe^2+^ cofactor in green; quantitative tunnel parameters are summarized in Table S4. **(c-d)** CASTpFold analysis of the putative hexameric interface pocket (yellow), dimeric interface pockets (red) and catalytic pockets (blue) of HGD-WT. Comparative analysis across variants revealed that the solvent accessible **(e)** surface area (SA) and **(f)** volume (SV) of the hexameric interface pocket decreases when introducing S54D or D86H AA substitutions whereas no changes were observed in **(g)** SA or **(h)** SV for the three calculated dimeric interface pockets. Interestingly, significantly decreased **(i)** SA and **(j)** SV for each of the six catalytic pockets could be observed for one high-activity variant containing the double mutation Q354P+P359E.

In addition to substrate access, catalytic performance in oxygenases such as HGD may also be influenced by dioxygen diffusion, which is thought to occur through transient cavities by thermal fluctuations rather than permanent channels. Previous studies have shown that HGD dynamics promotes the formation of transient internal pockets connecting the central opening of the hexamer to the active site [56,57]. Although oxygen diffusion pathways were not explicitly analysed in the present study, they may nonetheless be relevant, as position 401 was included in the SSM and has previously been reported as a key residue involved in the dioxygen diffusion pathway rather than in structural packing. Notably, no beneficial substitutions were identified at this position, which is consistent with earlier studies showing that even conservative replacements at residue 401 can lead to AKU development, underscoring the strong functional constraints imposed on residues involved in dioxygen transport. Future studies explicitly addressing oxygen diffusion using molecular dynamics-based simulation approaches, such as implicit ligand sampling, may further refine the identification of residues involved in the dioxygen gas diffusion pathway of human HGD [4,57,58].

However, when considering the complexity of engineering multimeric enzymes such as HGD, the observed activity improvements are notable. Unlike monomeric proteins, mutations in multimeric systems can simultaneously affect subunit stability and oligomer assembly, while protein-protein interfaces remain highly constrained and difficult to predict, and are therefore rarely explicitly targeted in engineering workflows [53,59–64]. In this context, the KnowVolution campaign identified five beneficial substitutions (Asn31, Ser54, Asp86, Gln354, Pro359) located in surface, interface, or core regions of the HGD hexamer rather than within the catalytic pocket itself. This suggests that the observed activity increases arise from indirect effects on structural stability, local dynamics, or subunit organization, consistent with previous reports on multimeric enzymes where subtle stabilization of quaternary structure enhances catalytic performance [55,65–67]. Accordingly, these findings align with the activity-stability trade-off governing enzyme evolution, where distal mutations can modulate catalytic efficiency by influencing protein dynamics or stabilizing functionally relevant conformations [68,69]. More specifically, in the context of AKU, the modest magnitude of the observed activity gains likely reflects the high degree of evolutionary conservation of the HGD enzyme, which imposes strong functional constraints and renders many missense substitutions deleterious, as previously reported [3,4].

At the same time, these results reflect the challenges of navigating the protein fitness landscape: beneficial mutations are rare, most substitutions are neutral or deleterious, and sequence space near the active site is sparsely populated [70–72]. These constraints also explain why approaches like epPCR libraries, although easy to generate and capable of sampling mutations across the entire protein, often yield only modest improvements: most substitutions occur at residues distant from the active site, where effects on catalysis are limited. Mutations in catalytically relevant positions are comparatively rare, and the polymerases used for epPCR further introduce a bias toward transitions over transversions, further reducing library diversity and the likelihood of identifying variants with pronounced functional changes [73]. To overcome these limitations, further rounds of the KnowVolution strategy could be applied using our best variants as templates, while multi-site saturation mutagenesis approaches, such as OmniChange, allow simultaneous exploration of multiple positions and cooperative effects that might otherwise be missed [27,74].

However, further studies in physiologically relevant models will be necessary to verify that the experimentally observed catalytic activities in *E. coli* are reproducible under human-relevant conditions, first *in vitro* using human liver cell models expressing the improved HGD variants, and subsequently *in vivo* in an AKU mouse model [75]. Altogether, by linking sequence variants to functional outcomes, this strategy provides new insights into how specific AA changes affect HGD stability and activity, improves our understanding of genotype-phenotype correlations in AKU, and lays a foundation for potential precision medicine interventions.

## 4 Conclusions

In this study, we applied the KnowVolution strategy to systematically explore structure-function relationships in human HGD, an enzyme whose deficiency underlies AKU. By combining random and targeted mutagenesis with computational analysis and rational recombination, we identified amino acid positions that modulate catalytic activity without directly affecting the catalytic pocket. Across all phases of the workflow, beneficial substitutions were consistently located in surface-exposed or structural regions, highlighting the importance of indirect mechanisms in regulating HGD function.

Structural mapping and interface analysis indicated that certain positions, such as Asn31 and Asp86, contribute directly to hexamer stabilization, while structural inspection additionally suggested a role for Ser54 in intersubunit interactions at the dimeric interface. In contrast, substitutions at Gln354 and Pro359 likely influence activity through subtle effects on local structural dynamics in a region surrounding the Gly347-Gly355 loop, which acts as a dynamic lid over the active-site pocket. These findings suggest that distal and interface residues play an important role in modulating catalytic performance in HGD and may represent a broader design principle for multimeric enzymes.

Although the observed activity improvements are moderate, they underscore the complexity of engineering multimeric enzymes, where mutations can influence both catalytic function and oligomer stability. At the same time, the identification of beneficial substitutions outside the catalytic pocket emphasizes the functional relevance of regions that are often overlooked in protein engineering.

Overall, this work demonstrates that HGD activity can be modulated through distal structural elements and provides new insight into how non-catalytic residues contribute to enzyme function. In the context of AKU, these findings improve our understanding of genotype-phenotype relationships and offer a foundation for future studies aimed at evaluating HGD variants in physiologically relevant models and, ultimately, for informing the development of restorative therapeutic strategies.

## Acknowledgements

This study is financially supported by the Research Foundation Flanders (FWO) under grant number G023320N (EvolvAKUre). Figure 2 was created with BioRender.com (https://BioRender.com/6blr5jg).

## Declaration of generative AI and AI-assisted technologies in the writing process

During the preparation of this work, the authors used ChatGPT in order to improve the clarity and readability of certain sentences by refining grammar and restructuring complex phrases. After using this tool, the authors reviewed and edited the content as needed and take full responsibility for the content of the published article.

## CRediT authorship contribution statement

Conceptualization, S.L. and J.D.K.; Investigation, S.L., J.N., L.D., N.S.S., J.D.K; Methodology, S.L., J.D.K.; Resources, T.V. and J.D.K.; Data Curation, S.L. and J.D.K.; Writing – Original Draft Preparation, S.L. and J.D.K.; Writing – Review and Editing, S.L., J.N., L.D., N.S.S., T.V., U.S. and J.D.K; Visualization, S.L. and J.D.K.; Supervision, T.V., U.S., and J.D.K. Funding Acquisition, T.V. and J.D.K. All authors have read and approved the final manuscript.

## Additional Information

### Competing Interest Statement

S.L., J.D.K., J.N. and T.V. are named inventors on a pending patent application filed by the Vrije Universiteit Brussel (VUB) relating to engineered HGD variants with altered catalytic activity, aspects of which are described in this manuscript. The remaining authors declare that they have no known competing financial interests or personal relationships that could have appeared to influence the work reported in this paper.

### Data Availability Statement

The data generated or analysed during this study are included in this published article (and its Supplementary information files). Tunnel analyses were performed using CAVER Web. Full CAVER Web outputs are available under the following job IDs: k75ozm (HGD-WT), tymtde (HGD N31S+S54D+D86H), xtlfam (HGD D86H+Q354H), s5dl0v (HGD P359E+Q354P), dc6dkg (HGD S54D+D86H), haeqld (D86Q+Q354H). Pocket analyses were performed using CASTpFold. Full CASTpFold outputs, including complete residue lists, are available at j_69679addddfe0 (HGD-WT), j_6967a50c0b8de (HGD N31S+S54D+D86H), j_6967a7f4bd51e (HGD D86H+Q354H), j_6967a8f941261 (HGD P359E+Q354P), j_6967aa3b26f9c (HGD S54D+D86H), j_6967abc31cc88 (HGD D86Q+Q354H).

