## Supplementary Data for "Deciphering and Improving Human Homogentisate 1,2-Dioxygenase Function Through Knowledge Gaining Directed Evolution: Implications for Alkaptonuria"

**Supplementary Material**

**Supplementary Data 1.**

**HGD cDNA codon-optimized for *E. coli*:**

ATGGCCGAGCTGAAATATATCAGCGGTTTTGGTAATGAATGCAGCAGCGAAGATCCGCGTTGTCCGGGTAGCCTGCCGGAAGGTCAGAATAATCCGCAGGTTTGTCCGTATAATCTGTATGCAGAACAGCTGAGCGGTAGCGCATTTACCTGTCCGCGTAGCACCAATAAACGTAGCTGGCTGTATCGTATTCTGCCGAGCGTTAGCCATAAACCGTTTGAAAGCATTGATGAAGGTCAGGTTACCCATAATTGGGATGAAGTTGATCCTGATCCGAATCAGCTGCGTTGGAAACCTTTTGAAATTCCGAAAGCAAGCCAGAAAAAAGTGGATTTTGTTAGCGGTCTGCATACCCTGTGTGGTGCCGGTGATATCAAAAGCAATAATGGTCTGGCCATCCATATCTTTCTGTGTAATACCAGCATGGAAAATCGCTGCTTTTATAACAGTGATGGCGATTTTCTGATTGTGCCGCAGAAAGGTAATCTGCTGATTTATACCGAATTTGGCAAAATGCTGGTTCAGCCGAATGAAATTTGTGTTATTCAGCGTGGTATGCGCTTTAGCATCGATGTTTTTGAAGAAACCCGTGGCTATATCCTGGAAGTTTATGGTGTTCATTTTGAACTGCCGGATCTGGGTCCGATTGGTGCAAATGGCCTGGCAAATCCGCGTGATTTCCTGATTCCGATTGCATGGTATGAAGATCGTCAGGTTCCGGGTGGTTATACCGTTATTAACAAATATCAGGGCAAACTGTTTGCAGCCAAACAGGATGTTAGCCCGTTTAATGTTGTTGCATGGCATGGTAATTATACCCCGTATAAATACAACCTGAAAAACTTCATGGTGATCAACAGCGTTGCATTTGATCATGCAGATCCGAGCATTTTTACCGTTCTGACCGCAAAAAGCGTGCGTCCGGGTGTTGCAATTGCCGATTTTGTGATTTTTCCGCCTCGTTGGGGTGTTGCCGATAAAACCTTTCGTCCGCCTTATTATCATCGTAATTGCATGAGCGAGTTTATGGGTCTGATTCGTGGTCATTATGAAGCAAAACAGGGTGGTTTTCTGCCTGGTGGTGGTAGTCTGCATAGCACCATGACACCGCATGGTCCGGATGCAGATTGTTTTGAAAAAGCCAGCAAAGTTAAACTGGCACCGGAACGTATTGCAGATGGCACCATGGCATTTATGTTTGAAAGTAGCCTGAGCCTGGCAGTTACCAAATGGGGTCTGAAAGCCAGCCGTTGTCTGGATGAAAATTATCATAAATGTTGGGAGCCGCTGAAAAGCCATTTTACCCCGAATAGCCGTAATCCGGCAGAACCGAATTAA

**Supplementary Data 2.**

**Custom phosphorothioate (PTO) primers used for PCR amplification of HGD insert and pET42b backbone for PLICing:**

*FW Insert (HGD)*

5'- ctt taa gaa gga gat ata caT ATG GCC GAG CTG -3'

*RV Insert (HGD):*

5'- tta gca gcc tag gtA TTA ATC AAT TAG TGG T -3'

*FW Vector (pET42b)*

5'- acc tag gct gct aaA CAA AGC -3'

*RV Vector (pET42b):*

5'- tgt ata tct cct tct taa agT TAA ACA AAA TTA TTT CTA G -3'

**Supplementary Methods S1. Assay optimization**

**S1.1 Kinetic evaluation of enzyme-to-assay mix ratios**

The kinetic evaluation of enzyme-to-assay mix ratios was performed using the high-throughput HGD activity assay previously published by Lequeue *et al.* [1], with protein expression and cell lysis carried out in a 96-well deep-well plate format as detailed in the main article (Section 2.4.1.1, Primary library screening). Cleared lysates of *E. coli* BL21 (DE3) expressing HGD-WT or EV controls were assayed at enzyme-to-assay mix ratios of 10:90, 15:85, 20:80, and 25:75 (*v/v*) to a final volume of 100 µL. For each ratio, eight technical replicates were performed for both HGD-WT and EV, and the experiment was repeated in duplicate (two independent biological replicates). HGD enzymatic activity was measured, and enzyme velocities were calculated exactly as described in our previously published method [1]. Absorbance at 340 nm was recorded every 30 s for 25 min at 37 °C to generate reaction progress curves, which were subsequently used to determine the optimal enzyme-to-assay mix ratio and the linear reaction range. Because enzyme linearity had already been established across a range of enzyme-to-assay mix ratios under the primary screening conditions, re-evaluation for the re-screen was limited to a targeted assessment of the 10:90 ratio. For this purpose, linearity under re-screen conditions was evaluated in a single biological run, following the protocol described in detail in Section 2.4.1.2.

**S1.2 Dose-response analysis for selection of the optimal HGA concentration**

To determine the HGA concentration at which the reaction reaches its maximum velocity (V_MAX_), a dose-response experiment was performed using the previously optimized enzyme-to-assay mix ratio of 25/75 for the primary screen (*v/v*). HGA concentrations ranging from 0 to 3 mM were tested (0, 0.2, 0.4, 0.5, 0.6, 0.7, 0.8, 0.9, 1, 1.5, 2, and 3 mM), following the protocol described in Section 2.4.1.1. Each HGA concentration was tested in six technical replicates for the HGD-WT enzyme, while the EV negative control was included in two technical replicates. The experiment was repeated in three independent biological batches. Absorbance at 340 nm was recorded every 30 s for 25 min at 37 °C, a duration chosen to remain within the linear reaction range, and initial reaction velocities were calculated as described previously [1].

**Supplementary Methods S2. Assay robustness**

**S2.1 Plate uniformity, drift, and edge-effect analysis**

Assay robustness was assessed using a plate uniformity study as described previously [1], with HGD_G161R_ (low signal), HGD_E42A_ (medium signal), and HGD-WT (high signal) serving as benchmarks, with the only modification being that protein expression was carried out in 96-well plates rather than Erlenmeyer flasks. Each condition was tested in triplicate over three independent days. All experiments in this section were performed as described under the final optimized conditions (see Section 2.4.3), including culture, induction, lysis, and enzyme activity measurements. Enzyme velocities and statistical parameters (mean, standard deviation (SD), coefficient of variation (CV), signal window (SW), and Z’ factor) were calculated according to the previously published method [1]. Drift and edge effects were evaluated by comparing column-wise and row-wise signal distributions to the plate median. Column-wise and row-wise drift were assessed by calculating the relative deviation of median signal intensities along horizontal (left-to-right) and vertical (top-to-bottom) axes, respectively. Edge effects were calculated as the relative difference between median signal intensities of edge wells and interior wells. All positional effects are reported as percentage deviations relative to the plate median.

**Supplementary Results S1. Assay Optimization.**

**S1.1 Enzyme-to-assay mix ratio optimization**

Reliable quantification of enzymatic activity requires careful optimization of enzyme concentration to ensure that reaction rates are measured within the linear phase of the progress curve, corresponding to proportional substrate turnover or product accumulation. Under substrate-saturating conditions, where the reaction operates at its maximum velocity (V_MAX_), the observed reaction rate scales linearly with enzyme concentration, allowing accurate quantification of catalytic activity [2]. In the initial experiments, reaction progress curves were recorded under the primary screening conditions to identify an appropriate ratio of cell lysate containing HGD-WT and assay mix containing HGA. Enzyme-to-assay mix ratios of 10:90, 15:85, 20:80, and 25:75 (*v/v*) were tested at a final HGA concentration of 2.5 mM. At this substrate concentration, the HGD active sites are expected to be saturated, allowing the reaction to proceed at maximal velocity and independently of substrate concentration **(Fig. S1a)**. Linear regression analysis demonstrated significant linearity for all ratios over the 0-25 min interval (p < 0.0001). The goodness of fit increased with higher enzyme content, with R^2^ values of 0.80, 0.83, 0.90, and 0.94 for the 10:90, 15:85, 20:80, and 25:75 ratios, respectively. The strongest linearity was observed at the 25:75 enzyme-to-assay ratio between 1.3 and 21.3 min (R^2^ = 0.98; **Fig. S1b**). These results established the 25:75 ratio as suitable for reliable quantification of enzyme activity under primary screening conditions using the slope of the reaction progress curve (∆Abs_MAA_/minute). Following this primary optimization, a re-screen format incorporating minor procedural changes was implemented. Because enzyme linearity had already been established across a range of ratios, re-evaluation was limited to a targeted assessment of the 10:90 enzyme-to-assay mix ratio. Under re-screen conditions, linearity at the 10:90 enzyme-to-assay ratio was evaluated in a single biological run between 1.3 and 18.7 min (R^2^ = 0.99; **Fig. S1b**) and subsequently confirmed during robustness testing. Notably, the close overlap of the reaction progress curves obtained at the 10:90 and 25:75 enzyme-to-assay mix ratios indicates that both conditions operate within the same linear kinetic regime, such that further evaluation of additional ratios was not required for the re-screen.

**S1.2 Substrate concentration optimization**

Although previously published Michaelis-Menten parameters were obtained using protein expressed in Erlenmeyer flasks, the substrate concentration was re-optimized following the transition to a 96-well expression format. Under the primary screening conditions, a substrate titration was performed at the optimized enzyme-to-assay mix ratio of 25:75 (*v/v*) to identify an HGA concentration that supports near-V_MAX_ conditions. HGA concentrations from 0 to 3 mM (0, 0.2, 0.4, 0.5, 0.6, 0.7, 0.8, 0.9, 1, 1.5, 2, and 3 mM) were evaluated. Initial rates increased with HGA concentration up to approximately 2 mM and then approached a plateau across three batches (**Fig. S1c**), with only small and batch-dependent changes between 2 and 3 mM. Accordingly, 2.5 mM HGA was selected as a near-saturating substrate concentration for subsequent experiments. For the re-screen, substrate titration experiments were not repeated, and 2.5 mM HGA was used throughout to maintain consistency with the primary screening conditions.

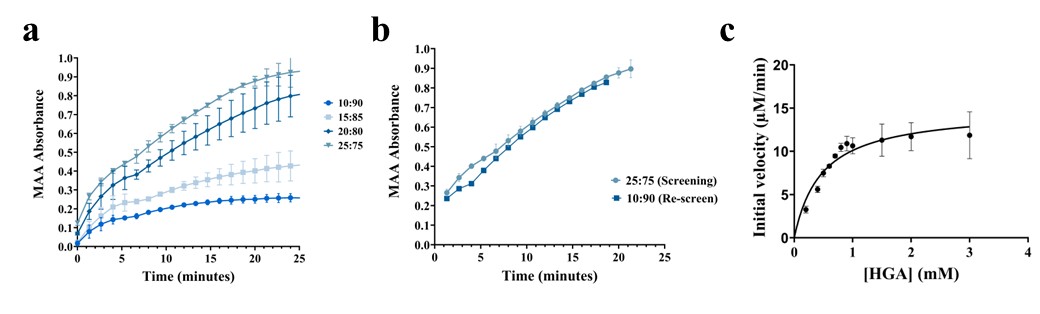

**Figure S1. Optimization of assay conditions for reliable quantification of HGD activity. (a)** Reaction progress curves for HGD-WT measured under primary screening conditions using enzyme-to-assay mix ratios of 10:90, 15:85, 20:80, and 25:75 (*v/v*) at a final HGA concentration of 2.5 mM. Absorbance corresponding to MAA formation was monitored over 25 min. Linear regression analysis demonstrated significant linearity for all ratios over the 0-25 min interval (p < 0.0001), with increasing goodness of fit at higher enzyme content (R^2^ = 0.80, 0.83, 0.90, and 0.94 for the 10:90, 15:85, 20:80, and 25:75 ratios, respectively). **(b)** Linear regions of the reaction progress curves for the 25:75 ratio under primary screening conditions and the 10:90 ratio under re-screen conditions, demonstrating comparable linearity. **(c)** Substrate titration performed under primary screening conditions at the 25:75 ratio, showing saturation of initial rates at HGA concentrations ≥ 2 mM; 2.5 mM HGA was selected for subsequent experiments.

**Supplementary Results S2. Assay robustness**

***S2.1 Plate uniformity and signal variability assessment.***

Robustness testing was restricted to the re-screen assay, as this stage was designed for reliable quantitative evaluation of enzyme activity, whereas the primary screen served as an initial qualitative selection step. Assay robustness was evaluated essentially as described in our previously published protocol [1], with the sole modification that protein expression was performed in 96-well deep-well plates rather than Erlenmeyer flasks. Validation experiments were conducted over three independent days, with three plates processed per day to assess plate uniformity, signal separation, and variability under screening conditions [5]. Uniformity testing was performed using benchmark controls representing high, medium, and low HGD activity levels. The high signal was defined by HGD-WT, corresponding to 100% enzymatic activity and maximal MAA formation. The low signal was defined by the HGD_G161R_ variant, which exhibits <1% residual HGD activity according to the literature, while the medium signal was represented by the HGD_E42A_ variant, with approximately 29% residual activity [6,7]. These three control signals were systematically interleaved across each plate and rotated daily in predefined layouts (high-medium-low, low-high-medium, and medium-low-high) to enable detection of positional effects, including edge effects and signal drift. Signal distributions obtained from all plates across three consecutive experimental days are summarized in **Figure S2** to provide an overview of day-to-day consistency. Plate uniformity data obtained in the 96-well format are summarized in **Table S1** and were used to calculate standard assay performance metrics as previously described [1]. CVs were determined for each signal level, with acceptance criteria requiring CV values ≤ 20%. CVs for the high and medium signals were consistently below this threshold (1.3-4.7% and 2.6-11.0%, respectively), while CVs for the low signal occasionally exceeded 20%, reflecting the inherently low mean signal of the HGD_G161R_ control. As an alternative acceptance criterion, the standard deviation of the minimum signal (SD_min_) was required to be less than or equal to the standard deviations of the medium (SD_mid_) and maximum signals (SD_max_), which was satisfied across all plates. Overall assay performance was further supported by average Z′ factors and signal window (SW) values of 0.91 ± 0.05 and 45.04 ± 23.05, respectively, exceeding predefined acceptance thresholds (Z′ > 0.4; SW > 2). In addition, normalized medium-signal variability (mid%) remained below 20%, and both within-day and between-day fold shifts were <2. Together, these results demonstrate that miniaturization of protein expression to a 96-well format did not compromise assay robustness and that the assay reliably discriminates HGD variants based on catalytic activity under HTS conditions.

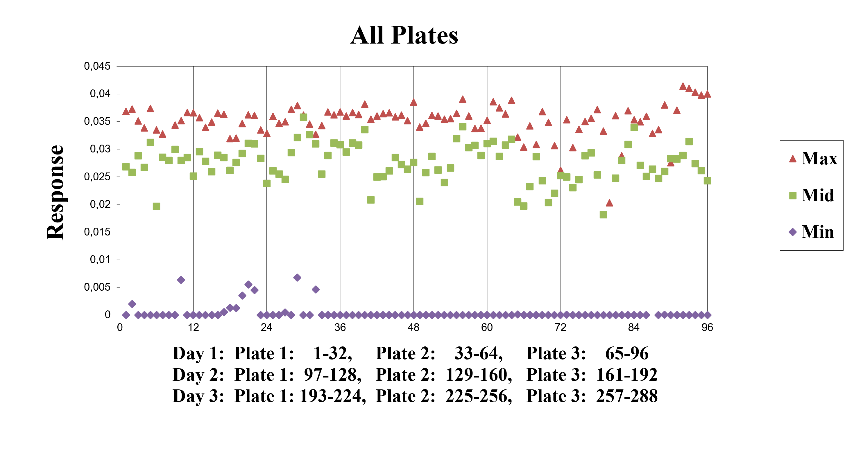

**Figure S2. Summary of control signal distributions across three experimental days.** Signal distributions obtained from all plates measured over three consecutive days are shown to provide an overview of day-to-day consistency. Max (HGD-WT, 100% activity), Mid (HGD_E42A_, ~72% residual activity), and Min (HGD_G161R_, <1% activity) control signals were systematically interleaved and rotated across plates to assess assay uniformity, signal separation, and positional effects.

***S2.2 Spatial uniformity assessment.***

According to the HTS Assay Validation Guidelines, a key requirement for HTS systems is the absence of substantial edge or drift effects, with signal variations below 20% considered negligible [5]. Scatter plots were used to visualize signal distribution and to identify potential patterns indicative of drift, edge effects, or other sources of systematic variability. Signals were plotted against well number in both column-wise and row-wise orientations (**Figure S3**). To mitigate potential edge effects arising from incubation-related factors such as temperature gradients, and to facilitate detection of systematic errors including drift, an interleaved plate layout was employed. Drift in Max and Mid signals was evaluated by analysing horizontal (left-to-right) and vertical (top-to-bottom) trends. For the Max signal, a column-wise drift above the 20% threshold was observed in one plate, while the same plate layout on other days did not exhibit this pattern. As the effect was not consistent across plates, it was not considered indicative of a systematic drift. Excluding this plate, column-wise drift ranged from 1.90% to 8.69%. In the row-wise analysis, elevated drift exceeding 20% was observed in two individual plates. However, this effect was not reproducible across plates generated using the same layout on other days. As the observed drift was not consistent, it was not considered indicative of a systematic row-wise effect. When these plates were excluded, row-wise drift values ranged from 3.94% to 19.15%. For the Mid signal, a reproducible column-wise drift was observed for plate 3 across all three experimental days using the same plate layout. The effect was consistently restricted to the first column and was characterized by elevated drift values of 48.17%, 27.98%, and 20.51% across the three experimental days. The stability of both the magnitude and spatial position of this effect indicates a systematic layout-associated bias rather than random variability. To prevent the introduction of systematic bias in subsequent experiments, the first column was excluded from Mid-signal sampling. In contrast, column-wise drift for the Mid signal in plates 1 and 2 was markedly lower, ranging from 5.63% to 15.85%, indicating that the pronounced drift was specific to plate 3 and not a general feature across all plates. Row-wise variability in the Mid signal exceeded 20% in several plates. However, no consistent directional trend (top-to-bottom or bottom-to-top) was observed. As the variation did not follow a systematic spatial pattern, it was not considered indicative of a true row-wise drift effect. After exclusion of plates with elevated variability, residual row-wise variation ranged from 13.78% to 18.90%. Edge effects for the Max signal were evaluated for both column-wise and row-wise orientations. Across all plates, edge-effect values remained low and did not exceed 17% in either direction. Column-wise edge effects ranged from -3.53% to 16.35%, while row-wise edge effects ranged from -13.07% to 17.00%, with negative values indicating lower signal intensities at plate edges relative to interior wells. No consistent enrichment or depletion of signal at plate edges was observed, indicating the absence of a systematic edge effect for the Max signal. Edge effects for the Mid signal were assessed in both column-wise and row-wise orientations. Apart from this layout-specific first-column effect, elevated edge-effect values were not reproducible across plates or experimental days and did not follow a consistent spatial pattern characteristic of classical edge effects. After exclusion of plates exhibiting edge-effect values above 20%, the remaining column-wise edge effects for the Mid signal ranged from -15.35% to 16.39%, while row-wise edge effects ranged from -4.42% to 11.41%. Collectively, these results indicate the absence of a significant or systematic global edge effect for the Mid signal.

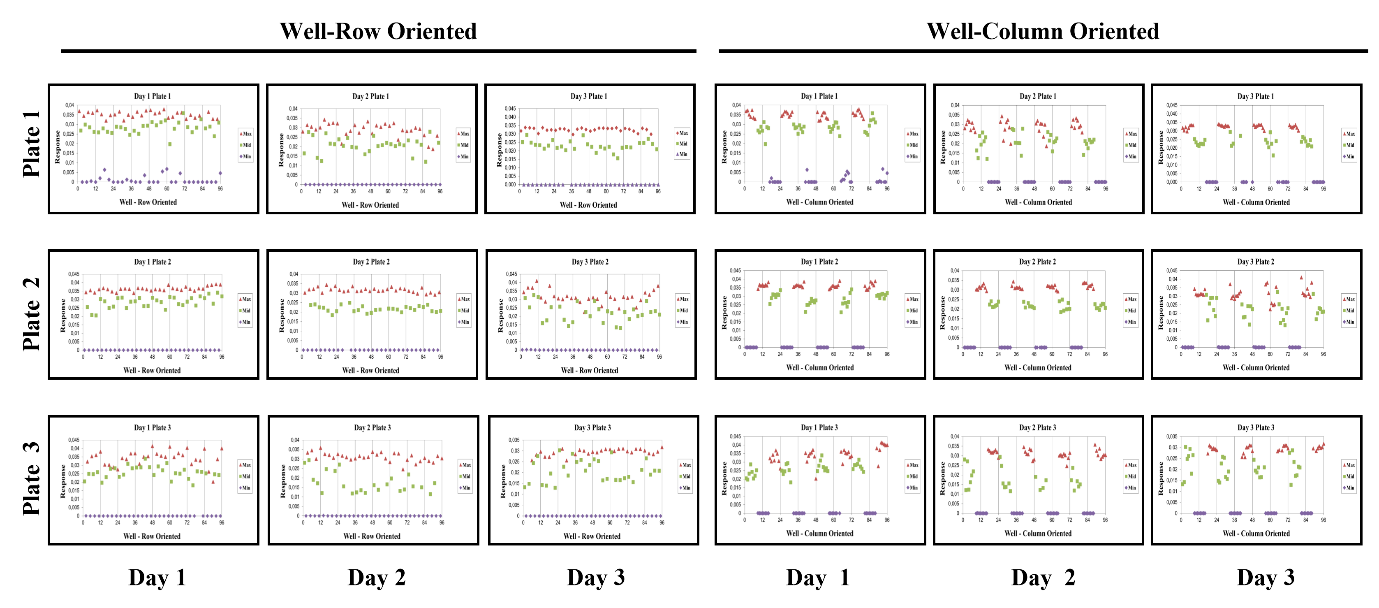
**Figure S3. Assessment of positional effects across plates and experimental days.** Scatter plots show signal intensities plotted against well number for individual plates measured over three consecutive days (three plates per day), displayed in column-wise and row-wise orientations to assess drift and edge effects. Max and Mid control signals were evaluated for horizontal (left-to-right) and vertical (top-to-bottom) trends. For most plates, no systematic drift or edge effects were observed. Isolated deviations exceeding 20% occurred in individual plates but were not reproducible across plates or days and were therefore not considered indicative of positional bias. In contrast, a reproducible column-wise effect affecting the Mid signal was observed in one plate layout and was mitigated by exclusion of the first column from subsequent analyses.

| Day | Plate | Type | Mean | SD | CV | Mid% | SW | Z’ |
| --- | --- | --- | --- | --- | --- | --- | --- | --- |
| 1 | 1 | H | 0.035 | 0.002 | 1.67 | 79.62 ± 8.86 | 51.00 | 0.88 |
|  |  | M | 0.028 | 0.003 | 3.77 |  |  |  |
|  |  | L | 0.001 | 0.002 | 64.6 |  |  |  |
|  | 2 | H | 0.036 | 0.001 | 1.35 | 78.46 ± 9.12 | 70.96 | 0.96 |
|  |  | M | 0.028 | 0.003 | 4.11 |  |  |  |
|  |  | L | 0.000 | 0.000 | N.D. |  |  |  |
|  | 3 | H | 0.034 | 0.005 | 4.73 | 75.26 ± 10.34 | 18.13 | 0.86 |
|  |  | M | 0.026 | 0.004 | 4.86 |  |  |  |
|  |  | L | 0.000 | 0.000 | N.D. |  |  |  |
| 2 | 1 | H | 0.029 | 0.004 | 4.58 | 72.32 ± 14.77 | 18.82 | 0.86 |
|  |  | M | 0.021 | 0.004 | 7.22 |  |  |  |
|  |  | L | 0.000 | 0.000 | N.D. |  |  |  |
|  | 2 | H | 0.032 | 0.001 | 1.45 | 68.79 ± 5.13 | 65.82 | 0.96 |
|  |  | M | 0.022 | 0.002 | 2.64 |  |  |  |
|  |  | L | 0.000 | 0.000 | N.D. |  |  |  |
|  | 3 | H | 0.031 | 0.002 | 2.59 | 56.66 ± 17.67 | 35.54 | 0.92 |
|  |  | M | 0.018 | 0.005 | 11.02 |  |  |  |
|  |  | L | 0.000 | 0.000 | N.D. |  |  |  |
| 3 | 1 | H | 0.032 | 0.001 | 1.28 | 72.00 ± 8.70 | 74.96 | 0.96 |
|  |  | M | 0.023 | 0.003 | 4.57 |  |  |  |
|  |  | L | 0.000 | 0.000 | N.D. |  |  |  |
|  | 2 | H | 0.032 | 0.004 | 4.53 | 70.37 ± 17.12 | 19.03 | 0.86 |
|  |  | M | 0.022 | 0.005 | 8.59 |  |  |  |
|  |  | L | 0.000 | 0.000 | N.D. |  |  |  |
|  | 3 | H | 0.029 | 0.002 | 1.85 | 70.65 ± 17.41 | 51.08 | 0.94 |
|  |  | M | 0.021 | 0.005 | 8.71 |  |  |  |
|  |  | L | 0.000 | 0.000 | N.D. |  |  |  |

**Table S1**. Data of the (intra)plate uniformity study. H: high signal; M: medium signal; L: low signal; SD: standard deviation; CV: coefficient of variation; Mid%: normalized mid signal; SW: signal window; Z’: Z’ factor. All maximum (high) and mid signal (unnormalized) CVs were <20%. All normalized mid signal (Mid%) SDs were <20%. All minimum (low) SDs were lower than the corresponding maximum (high) and mid SDs. All signal windows (SW) were >2 and all Z’ factors were >0.4. Z’ factors and CVs were calculated based on eight replicates per equivalent signal. Within-day and between-day fold shifts were below two, indicating robust assay performance. Based on these uniformity data, a reproducible first-column drift was identified for the Mid signal and was subsequently excluded from Mid-signal sampling in follow-up experiments.

**Table S2**. 22c-trick primers used for SSM at positions identified from Phase I of KnowVolution. For each targeted position, a primer pair was designed using the 22c-trick degenerate codon strategy to introduce the remaining 19 non-WT amino acids at the selected site in Phase II. Degenerate bases are according to the IUPAC code: N = A/C/G/T; D = A/G/T; V = A/C/G; H = A/C/T; B = C/G/T. For each primer, the primer ID, exact annealing temperature (Ta; °C) used in the PCR reaction, and sequence (5'→3') are provided.

| **Position** | **Primer ID** | **Sequence (5’ – 3’)** |  | **T_a_ (°C)** |
| --- | --- | --- | --- | --- |
| **31** | NDT (Fw) | GAAGGTCAGAATNDTCCGCAGGTTTG |  | 60 |
|  | AHN (Rv) | CAAACCTGCGGAHNATTCTGACCTTC |  |  |
|  | VHG (Fw) | GAAGGTCAGAATVHGCCGCAGGTTTG |  |  |
|  | CDB (Rv) | CAAACCTGCGGCDBATTCTGACCTTC |  |  |
|  | TGG (Fw) | GAAGGTCAGAATTGGCCGCAGGTTTG |  |  |
|  | CCA (Rv) | CAAACCTGCGGCCAATTCTGACCTTC |  |  |
| **317** | NDT (Fw) | GCCGATTTTGTGNDTTTTCCGCCTCGT |  | 60 |
|  | AHN (Rv) | ACGAGGCGGAAAAHNCACAAAATCGGC |  |  |
|  | VHG (Fw) | GCCGATTTTGTGVHGTTTCCGCCTCGT |  |  |
|  | CDB (Rv) | ACGAGGCGGAAACDBCACAAAATCGGC |  |  |
|  | TGG (Fw) | GCCGATTTTGTGTGGTTTCCGCCTCGT |  |  |
|  | CCA (Rv) | ACGAGGCGGAAACCACACAAAATCGGC |  |  |
| **223** | NDT (Fw) | AATGGCCTGGCANDTCCGCGTGATTTC |  | 72 |
|  | AHN (Rv) | GAAATCACGCGGAHNTGCCAGGCCATT |  |  |
|  | VHG (Fw) | AATGGCCTGGCAVHGCCGCGTGATTTC |  |  |
|  | CDB (Rv) | GAAATCACGCGGCDBTGCCAGGCCATT |  |  |
|  | TGG (Fw) | AATGGCCTGGCATGGCCGCGTGATTTC |  |  |
|  | CCA (Rv) | GAAATCACGCGGCCATGCCAGGCCATT |  |  |
| **54** | NDT (Fw) | ACCTGTCCGCGTNDTACCAATAAACGT |  | 69 |
|  | AHN (Rv) | ACGTTTATTGGTAHNACGCGGACAGGT |  |  |
|  | VHG (Fw) | ACCTGTCCGCGTVHGACCAATAAACGT |  |  |
|  | CDB (Rv) | ACGTTTATTGGTCDBACGCGGACAGGT |  |  |
|  | TGG (Fw) | ACCTGTCCGCGTTGGACCAATAAACGT |  |  |
|  | CCA (Rv) | ACGTTTATTGGTCCAACGCGGACAGGT |  |  |
| **86** | NDT (Fw) | ACCCATAATTGGNDTGAAGTTGATCCT |  | 65 |
|  | AHN (Rv) | AGGATCAACTTCAHNCCAATTATGGGT |  |  |
|  | VHG (Fw) | ACCCATAATTGGVHGGAAGTTGATCCT |  |  |
|  | CDB (Rv) | AGGATCAACTTCCDBCCAATTATGGGT |  |  |
|  | TGG (Fw) | ACCCATAATTGGTGGGAAGTTGATCCT |  |  |
|  | CCA (Rv) | AGGATCAACTTCCCACCAATTATGGGT |  |  |
| **91** | NDT (Fw) | GAAGTTGATCCTNDTCCGAATCAGCTG |  | 69 |
|  | AHN (Rv) | CAGCTGATTCGGAHNAGGATCAACTTC |  |  |
|  | VHG (Fw) | GAAGTTGATCCTVHGCCGAATCAGCTG |  |  |
|  | CDB (Rv) | CAGCTGATTCGGCDBAGGATCAACTTC |  |  |
|  | TGG (Fw) | GAAGTTGATCCTTGGCCGAATCAGCTG |  |  |
|  | CCA (Rv) | CAGCTGATTCGGCCAAGGATCAACTTC |  |  |
| **150** | NDT (Fw) | CGCTGCTTTTATAACNDTGATGGCGATTTTCTG |  | 70 |
|  | AHN (Rv) | CAGAAAATCGCCATCAHNGTTATAAAAGCAGCG |  |  |
|  | VHG (Fw) | CGCTGCTTTTATAACVHGGATGGCGATTTTCTG |  |  |
|  | CDB (Rv) | CAGAAAATCGCCATCCDBGTTATAAAAGCAGCG |  |  |
|  | TGG (Fw) | CGCTGCTTTTATAACTGGGATGGCGATTTTCTG |  |  |
|  | CCA (Rv) | CAGAAAATCGCCATCCCAGTTATAAAAGCAGCG |  |  |
| **354** | NDT (Fw) | TATGAAGCAAAANDTGGTGGTTTTCTG |  | 65 |
|  | AHN (Rv) | CAGAAAACCACCAHNTTTTGCTTCATA |  |  |
|  | VHG (Fw) | TATGAAGCAAAAVHGGGTGGTTTTCTG |  |  |
|  | CDB (Rv) | CAGAAAACCACCCDBTTTTGCTTCATA |  |  |
|  | TGG (Fw) | TATGAAGCAAAATGGGGTGGTTTTCTG |  |  |
|  | CCA (Rv) | CAGAAAACCACCCCATTTTGCTTCATA |  |  |
| **336** | NDT (Fw) | CCGCCTTATTATCATNDTAATTGCATGAGCGAG |  | 69 |
|  | AHN (Rv) | CTCGCTCATGCAATTAHNATGATAATAAGGCGG |  |  |
|  | VHG (Fw) | CCGCCTTATTATCATVHGAATTGCATGAGCGAG |  |  |
|  | CDB (Rv) | CTCGCTCATGCAATTCDBATGATAATAAGGCGG |  |  |
|  | TGG (Fw) | CCGCCTTATTATCATTGGAATTGCATGAGCGAG |  |  |
|  | CCA (Rv) | CTCGCTCATGCAATTCCAATGATAATAAGGCGG |  |  |
| **347** | NDT (Fw) | ATGGGTCTGATTNDTGGTCATTATGAA |  | 65 |
|  | AHN (Rv) | TTCATAATGACCAHNAATCAGACCCAT |  |  |
|  | VHG (Fw) | ATGGGTCTGATTVHGGGTCATTATGAA |  |  |
|  | CDB (Rv) | TTCATAATGACCCDBAATCAGACCCAT |  |  |
|  | TGG (Fw) | ATGGGTCTGATTTGGGGTCATTATGAA |  |  |
|  | CCA (Rv) | TTCATAATGACCCCAAATCAGACCCAT |  |  |
| **359** | NDT (Fw) | GGTGGTTTTCTGNDTGGTGGTGGTAGT |  | 71 |
|  | AHN (Rv) | ACTACCACCACCAHNCAGAAAACCACC |  |  |
|  | VHG (Fw) | GGTGGTTTTCTGVHGGGTGGTGGTAGT |  |  |
|  | CDB (Rv) | ACTACCACCACCCDBCAGAAAACCACC |  |  |
|  | TGG (Fw) | GGTGGTTTTCTGTGGGGTGGTGGTAGT |  |  |
|  | CCA (Rv) | ACTACCACCACCCCACAGAAAACCACC |  |  |
| **373** | NDT (Fw) | ACACCGCATGGTNDTGATGCAGATTGT |  | 71 |
|  | AHN (Rv) | ACAATCTGCATCAHNACCATGCGGTGT |  |  |
|  | VHG (Fw) | ACACCGCATGGTVHGGATGCAGATTGT |  |  |
|  | CDB (Rv) | ACAATCTGCATCCDBACCATGCGGTGT |  |  |
|  | TGG (Fw) | ACACCGCATGGTTGGGATGCAGATTGT |  |  |
|  | CCA (Rv) | ACAATCTGCATCCCAACCATGCGGTGT |  |  |
| **401** | NDT (Fw) | GCATTTATGTTTNDTAGTAGCCTGAGC |  | 63 |
|  | AHN (Rv) | GCTCAGGCTACTAHNAAACATAAATGC |  |  |
|  | VHG (Fw) | GCATTTATGTTTVHGAGTAGCCTGAGC |  |  |
|  | CDB (Rv) | GCTCAGGCTACTCDBAAACATAAATGC |  |  |
|  | TGG (Fw) | GCATTTATGTTTTGGAGTAGCCTGAGC |  |  |
|  | CCA (Rv) | GCTCAGGCTACTCCAAAACATAAATGC |  |  |

**Table S3. Primers designed for recombination introduced by SDM.** Primer length (nucleotides; nt), melting temperature (Tm; °C), and GC content (%) were calculated using the OligoAnalyzer tool, and the exact annealing temperature (T_a_; °C) applied in the PCR reaction is provided. The underlined codon indicates the site of the introduced substitution.

| Variant | Primer ID | Sequence (5’ – 3’) | Polymerase  DMSO (%)  T_a_ (°C) |
| --- | --- | --- | --- |
| N31S+S54D | Fw primer  S54D | CTGTCCGCGTGATACCAATAAAC | Phusion HF  1% DMSO  66 |
|  | Rv primer  S54D | GTTTATTGGTATCACGCGGACAG |  |
| P359M+Q354V | Fw primer  Q354V | GAAGCAAAAGTGGGTGGTTTTC | Phusion HF  1% DMSO  65 |
|  | Rv primer  Q354V | GAAAACCACCCACTTTTGCTTC |  |
| P359M+Q354H | Fw primer  Q354H | GAAGCAAAACATGGTGGTTTTC | Phusion HF  1% DMSO  63 |
|  | Rv primer  Q354H | GAAAACCACCATGTTTTGCTTC |  |
| P359E+Q354V | Fw primer  Q354V | GAAGCAAAAGTGGGTGGTTTTC | Phusion HF  1% DMSO  65 |
|  | Rv primer Q354V | GAAAACCACCCACTTTTGCTTC |  |
| P359E+Q354H | Fw primer Q354H | GAAGCAAAACATGGTGGTTTTC | Phusion HF  1% DMSO  63 |
|  | Rv primer Q354H | GAAAACCACCATGTTTTGCTTC |  |
| S54D+Q354V | Rv primer  Q354P | GCAAAAGTGGGTGGTTTTCTGCCTGGTGGTGGTAGTCTG | Phusion Flash  1% DMSO  70 |
|  | Rv primer Q354V | GAAAACCACCCACTTTTGCTTCATAATGACCACGAATCAGACCCA |  |
| Q354V+D86H | Fw primer D86H | ATTGGCATGAAGTTGATCCTGATCCGAATC | Phusion HF  1% DMSO  69 |
|  | Rv primer D86H | ACTTCATGCCAATTATGGGTAACCTGACCTTC |  |
| D86Q+Q354V | Fw primer Q354V | GCAAAAGTGGGTGGTTTTCTGCCTGGTGGTGGTAGTCTG | Phusion Flash  1% DMSO  69 |
|  | Rv primer Q354V | GAAAACCACCCACTTTTGCTTCATAATGACCACGAATCAGACCCA |  |
| S54D+Q354P | Fw primer Q354P | CAAAACCGGGTGGTTTTCTGCCTGGTGGTGGTAGTCTG | Phusion Flash  1% DMSO  70 |
|  | Rv primer Q354P | GAAAACCACCCGGTTTTGCTTCATAATGACCACGAATCAGACCCA |  |
| Q354P+D86H | Fw primer D86H | ATTGGCATGAAGTTGATCCTGATCCGAATC | Phusion HF  1% DMSO  70 |
|  | Rv primer D86H | ACTTCATGCCAATTATGGGTAACCTGACCTTC |  |
| D86Q+Q354P | Fw primer Q354P | CAAAACCGGGTGGTTTTCTGCCTGGTGGTGGTAGTCTG | Phusion Flash  1% DMSO  70 |
|  | Rv primer Q354P | GAAAACCACCCGGTTTTGCTTCATAATGACCACGAATCAGACCCA |  |
| Q354P+P359E | Fw primer Q354P+P359E | AACCGGGTGGTTTTCTGGAGGGTGGTGGTAGTCTGCATAGCACCATG | Phusion Flash  1% DMSO  70 |
|  | Rv primer  Q354P+P359E | CTCCAGAAAACCACCCGGTTTTGCTTCATAATGACCACGAATCAGACCCA |  |
| Q354P+P359M | Fw primer  Q354P+P359M | AACCGGGTGGTTTTCTGATGGGTGGTGGTAGTCTGCATAGCACCATG | Phusion Flash  1% DMSO  70 |
|  | Rv primer  Q354P+P359M | CATCAGAAAACCACCCGGTTTTGCTTCATAATGACCACGAATCAGACCCA |  |
| S54D+Q354H | Fw primer  Q354H | GCAAAACATGGTGGTTTTCTGCCTGGTGGTGGTAGTCTG | Phusion HF  1% DMSO  69 |
|  | Rv primer  Q354H | GAAAACCACCATGTTTTGCTTCATAATGACCACGAATCAGACCCA |  |
| Q354H+D86H | Fw primer  D86H | ATTGGCATGAAGTTGATCCTGATCCGAATC | Phusion HF  1% DMSO  70 |
|  | Rv primer  D86H | ACTTCATGCCAATTATGGGTAACCTGACCTTC |  |
| D86Q+Q354H | Fw primer  Q354H | GCAAAACATGGTGGTTTTCTGCCTGGTGGTGGTAGTCTG | Phusion Flash  70 |
|  | Rv primer  Q354H | GAAAACCACCATGTTTTGCTTCATAATGACCACGAATCAGACCCA |  |
| N31S+S54D+D86Q | Fw primer  D86Q | CATAATTGGCAGGAAGTTGATCCTGATCCGAATCAGCTGC | Phusion Flash  1% DMSO  69 |
|  | Rv primer  D86Q | CAACTTCCTGCCAATTATGGGTAACCTGACCTTCATCAATGC |  |
| S54D+D86Q | Fw primer  D86Q | CATAATTGGCAGGAAGTTGATCCTGATCCGAATCAGCTGC | Phusion Flash  1% DMSO  70 |
|  | Rv primer  D86Q | CAACTTCCTGCCAATTATGGGTAACCTGACCTTCATCAATGC |  |
| N31S+S54D+D86A | Fw primer  D86A | CATAATTGGGCTGAAGTTGATCCTGATCCGAATCAGCTGC | Phusion Flash  1% DMSO  67.3 |
|  | Rv primer  D86A | CAACTTCAGCCCAATTATGGGTAACCTGACCTTCATCAATGC |  |
| S54D+D86H | Fw primer  D86H | CATAATTGGCATGAAGTTGATCCTGATCCGAATCAGCTGC | Phusion Flash  1% DMSO  60.7 |
|  | Rv primer  D86H | CAACTTCATGCCAATTATGGGTAACCTGACCTTCATCAATGC |  |
| N31S+S54D+D86H | Fw primer  D86H | ATTGGCATGAAGTTGATCCTGATCCGAATC | Phusion HF  1% DMSO  70 |
|  | Rv primer  D86H | ACTTCATGCCAATTATGGGTAACCTGACCTTC |  |
| S54D+D86A | Fw primer  D86A | CATAATTGGGCTGAAGTTGATCCTGATCCGAATCAGCTGC | Phusion Flash  1% DMSO  70 |
|  | Rv primer  D86A | CAACTTCAGCCCAATTATGGGTAACCTGACCTTCATCAATGC |  |

**Table S4. Comparison of the primary substrate-access tunnel (tunnel 1) in HGD-WT and selected HGD mutants.** Tunnel 1, the main substrate-access pathway, was identified and analyzed using CAVER with identical parameters for all variants. Bottleneck radius (Å) is the narrowest point along the tunnel and is critical for substrate access. Length (Å) measures the distance from the active site to the protein surface, with shorter tunnels generally facilitating faster substrate entry. Throughput represents the CAVER priority score, integrating tunnel radius, length, and curvature. Key tunnel-lining residues are amino acids forming the bottlenecks that are directly and indirectly affected by mutations. Variants were selected to illustrate extremes of catalytic activity: the three highest-activity mutants, two nearly inactive mutants, and the WT. Note: Residues in parentheses indicate the chain from which they originate; all other residues are from the substrate-containing chain (chain B).

| Variant | Bottleneck radius (Å) | Length (Å) | Throughput | Key-lining residues |
| --- | --- | --- | --- | --- |
| WT | 1.8 | 19.4 | 0.7 | His292, Pro295, Phe298, Phe318, Arg321, Arg330, Pro331, Pro332, Tyr333, His335, Glu341, Met343, Tyr350, Glu351, Ala352, Lys353, His365, His371, Pro373, Cys377, Lys380, Ala381, Val384, Glu389, Ile391, Ala392, Met399, Phe49(D) |
| N31S+S54D+D86H (high) | 1.8 | 17.5 | 0.72 | His292, Pro295, Phe298, Phe318, Arg321, Arg330, Pro331, Pro332, Tyr333, His335, Glu341, Met343, Tyr350, Glu351, Ala352, Lys353, His365, His371, Pro373, Cys377, Lys380, Ala381, Val384, Glu389, Ile391, Ala392, Met399, Phe49(D) |
| D86H+Q354H (high) | 1.8 | 19.1 | 0.7 | His292, Pro295, Phe298, Phe318, Arg321, Arg330, Pro331, Pro332, Tyr333, His335, Glu341, Met343, Tyr350, Glu351, Ala352, Lys353, His365, His371, Pro373, Cys377, Lys380, Ala381, Val384, Glu389, Ile391, Ala392, Met399, Phe49(D) |
| P359E+Q354P (high) | 1.8 | 18.5 | 0.71 | His292, Pro295, Phe298, Phe318, Arg321, Arg330, Pro331, Pro332, Tyr333, His335, Glu341, Met343, Tyr350, Glu351, Ala352, Lys353, His365, His371, Pro373, Cys377, Lys380, Ala381, Val384, Glu389, Ile391, Ala392, Met399, Phe49(D) |
| S54D+D86H (low) | 1.8 | 17.1 | 0.72 | His292, Pro295, Phe298, Phe318, Arg321, Arg330, Pro331, Pro332, Tyr333, His335, Glu341, Met343, Tyr350, Glu351, Ala352, Lys353, His365, His371, Pro373, Cys377, Lys380, Ala381, Val384, Glu389, Ile391, Ala392, Met399, Phe49(D) |
| D86Q+Q354H (low) | 1.8 | 19.1 | 0.7 | His292, Pro295, Phe298, Phe318, Arg321, Arg330, Pro331, Pro332, Tyr333, His335, Glu341, Met343, Tyr350, Glu351, Ala352, Lys353, His365, His371, Pro373, Cys377, Lys380, Ala381, Val384, Glu389, Ile391, Ala392, Met399, Phe49(D) |
